# Complex sporulation-specific expression of transcription termination factor Rho highlights its involvement in *Bacillus subtilis* cell differentiation

**DOI:** 10.1101/2023.12.01.569620

**Authors:** Vladimir Bidnenko, Arnaud Chastanet, Christine Péchoux, Yulia Redko-Hamel, Olivier Pellegrini, Sylvain Durand, Ciarán Condon, Marc Boudvillain, Matthieu Jules, Elena Bidnenko

## Abstract

Transcription termination factor Rho controls pervasive, mainly antisense, transcription initiated at cryptic signals or resulting from read-through at weak terminators in various bacterial species. In *Bacillus subtilis*, Rho is intricately involved in the regulation of phenomena associated with the adaptation to stationary phase and cell differentiation including the ultimate survival program of sporulation.

While knockout or overexpression of the *rho* gene alters global transcription and modifies cell physiology, in wild-type *B. subtilis* cells, the reduction of Rho levels during the transition to stationary phase is necessary for both initiation and implementation of the sporulation program. However, the mechanisms that govern Rho expression throughout the cell cycle remain largely unknown.

Here, we demonstrate that, besides the previously identified vegetative SigA-dependent promoter active during exponential growth, two distinct mechanisms ensure a spatiotemporal expression of the *rho* gene during sporulation. In the mother cell of the sporangium, *rho* expression occurs through the read-through transcription initiated at the distal SigH-dependent and Spo0A∼P-regulated promoter of the *spo0F* gene. In the forespore, *rho* is transcribed from a genuine promoter recognized by the alternative sigma factor SigF. These regulatory elements compensate for the inactivation of SigA-dependent *rho* expression at the end of exponential growth and allow the critical “refueling” of Rho protein in both compartments of the sporangium. We show that altering *rho* expression in the mother cell or in the forespore affects differently the properties and the morphology of mature spores. Moreover, spores formed in the absence of Rho are impaired in their ability to revive under favorable growth conditions, exhibiting accelerated germination and slow outgrowth. Finally, we show that optimal outgrowth of the wild-type spores requires the expression of *rho* during spore maturation and additionally after spore germination.

## Introduction

A growing understanding of the importance of transcription termination in the regulation of gene expression in bacteria has stimulated extensive analysis of the proteins that control this universal step in decoding genetic information (Peters et al., 2011; Ray-Soni et al., 2016; Kriner et al., 2016; Turnbough, 2019; Mandell et al., 2022a). Among them is transcription termination factor Rho, an ATP-dependent RNA helicase-translocase that improves the termination efficiency of many intrinsic terminators and is essential for termination at bona-fide Rho-dependent terminators (Roberts, 1969; Quirk et al., 1996; Peters et al., 2009; Hao et al., 2021; Mandell et al., 2022b). Over the last years, considerable advances have been made in unveiling the molecular mechanism of Rho-dependent termination (Song et al., 2022; Molodtsov et al., 2023; Murayama et al., 2023; Rashid and Berger, 2023).

Transcriptome analyses of *rho* knockout and conditionally deficient mutants of various bacteria have established the essential role of Rho in controlling pervasive, mainly non-coding and antisense, transcription that originates from cryptic initiation signals or from read-through of transcription terminators (Nicolas et al., 2012, Peters et al., 2012; Mäder et al., 2016; Botella et al, 2017). By controlling transcription genome-wide, Rho directly and indirectly influences various aspects of cellular physiology in bacteria living in diverse habitats (Bidnenko et el., 2017, 2023; Botella et al., 2017; Nagel et al., 2018; Trzilova et al., 2020; Lin et al.; 2021; Krypotou et al., 2023). It is notable that Rho is involved in the regulation of phenomena associated with stationary phase, including stress survival, cell fate determination, antibiotic sensitivity, host colonization and virulence in different bacteria (Lee and Helmann, 2014; Liu et al., 2016; Hafeezunnisa et al., Bidnenko et al., 2017, 2023; Nagel et al., 2018; Trzilova et al. 2020; Lin et al., 2021; Figueroa-Bossi et al., 2022).

Recently, we provided evidence that Rho is intricately involved in the control of cellular adaptation to stationary phase and in cell-fate decision-making, in particular, the decision to sporulate, in the Gram-positive bacterium *Bacillus subtilis* (Bidnenko et al., 2017, 2023). The involvement of Rho in the sporulation process was also demonstrated in *Clostridioides difficile* and *Bacillus thuringiensis* (Trzilova et al., 2020; Lin et al.; 2021).

Sporulation is a complex developmental program that transforms a vegetative bacterial cell into a highly resistant dormant spore, thus ensuring the survival of bacteria under adverse environmental conditions and their dissemination within different ecological niches (de Hoon et al., 2010; Swick et al., 2016; Galperin et al., 2022). A hallmark of sporulation in *B. subtilis* is the asymmetric division of a differentiating cell in two unequal parts: a forespore, which further develops into the spore, and a mother cell, which engulfs the forespore, nourishes it, ensures the synthesis of spore protective layers, and finally lyses to release the mature spore. Proper spore morphogenesis is essential for spore resistance to external damages and its conversion back into a growing cell under favorable conditions through sequential processes of germination and outgrowth (Setlow, 2003; Setlow et al., 2017; Setlow and Christie, 2023; Segev et al., 2013; Abhyankar et al., 2016; Boone and Driks, 2016; Mutlu et al., 2018; 2020). In *B. subtilis*, sporulation is primarily controlled at the level of transcription initiation by the master regulator Spo0A, whose activity depends on phosphorylation mediated by a multi-component phosphorelay, the transition phase-specific sigma factor SigH and a cascade of sigma factors (SigF, SigE, SigG and SigK) that sequentially drive temporally- and spatially-defined transcriptional programs of spore morphogenesis. Sporulation is initiated at a threshold level of Spo0A∼P, which activates the expression of *sigH* and several sporulation genes, including *sigF* and *sigE* encoding the inactive forms of sigma factors. Upon completion of the asymmetric septum, activation of SigF in the forespore is required for subsequent activation of SigE in the mother cell. After a complete engulfment of the forespore, the compartment-specific programs of gene expression are further continued by SigG in the forespore and SigK in the mother cell (for comprehensives reviews see Errington, 1993; Stragier and Losick, 1996; Piggot and Hilbert, 2004; Higgins and Dworkin, 2012). The relevant regulons of each sporulation-specific sigma factor, as well as other genes involved in sporulation, were determined using the combination of experimental and bioinformatics approaches (Eichenberger et al, 2003, 2004; Steil et al., 2005; Wang et al., 2006; de Hoon et al., 2010; Overkamp et al., 2015; Meeske et al., 2016; Galperin et al., 2022).

We have previously shown that deletion of the *rho* gene in *B. subtilis* prevents intragenic termination of the *kinB* transcript, which encodes the sensor kinase KinB, one of the main kinases feeding phosphate into the Spo0A phosphorelay. This leads to the increased expression of KinB and, consequently, to rapid accumulation of active Spo0A∼P to a threshold level that triggers sporulation. Thus, Rho inactivation stimulates sporulation in *B. subtilis* indicating a regulatory role of Rho-mediated transcription termination within the cell fate decision-making network centered on Spo0A∼P (Bidnenko et al., 2017). Conversely, maintaining *rho* expression at a stably elevated level throughout *B. subtilis* exponential growth and stationary phase alters the early steps of adaptive reprogramming of cellular transcription, prevents the activation of Spo0A and, additionally, inhibits some late, yet non-identified, sporulation events, with the overall result of blocking the formation of spores (Bidnenko et al., 2023).

Comparative transcriptome and proteome analyses have revealed a decrease in the levels of *rho* mRNA and Rho protein during the transition to stationary phase in wild-type *B. subtilis* cells (Nicolas et al., 2012; Bidnenko et al., 2017; 2023). Considering that Rho negatively affected sporulation in our analyses, a decrease in Rho levels appears necessary for both initiation and implementation of the sporulation program. This in turn suggests that *rho* expression during *B. subtilis* growth and stationary phase is subject to reliable and timely regulation.

The only mechanism known to date for regulating *rho* expression is transcriptional attenuation at the Rho-dependent terminator(s) located within the leader region of the *rho* transcript and consequent premature transcription termination (Ingham et al., 1999). Similar to *B. subtilis,* transcription of the *rho* gene was shown to be autogenously regulated in *Escherichia coli*, *Salmonella* and *Caulobacter crescentus* (Barik et al., 1985; Matsumoto et al., 1986; Italiani et al., 2005; Silva et al., 2019). In *Salmonella*, *rho* autoregulation is counteracted by the small noncoding RNA SraL, which prevents the Rho-mediated termination by direct binding the 5′-UTR of *rho* mRNA (Silva et al., 2019).

In the present study, we aimed to gain further insights into *rho* regulation in *B. subtilis* by analyzing the kinetics of *rho* expression at various stages of cellular growth and differentiation. Quite unexpectedly, we found that the expression of the *rho* gene is specifically induced early during sporulation. This counterintuitive finding has focused our subsequent analysis on understanding the mechanism of the sporulation-specific expression of *rho* and its biological significance.

We describe two distinct mechanisms controlling *rho* expression in each compartment of the sporulating cells: read-through transcription from the upstream SigH-dependent promoter in the mother cell, and a genuine SigF-dependent *rho* promoter active in the forespore. The latter feature allows us to classify *rho* as a novel member of the SigF regulon of *B. subtilis*. We provide evidence that altering the spatiotemporal expression of *rho* affects spore resistance properties and morphology, in particular, the structure of spore coat. Moreover, spores formed in the absence of Rho are impaired in their ability to revive under favorable growth conditions, exhibiting an accelerated germination and a slow outgrowth. Finally, we show that the optimal rate of spore outgrowth depends on the synthesis of Rho during spore formation and *de novo* after germination.

## Results

### Rho expression is specifically induced during sporulation

To analyze a real-time expression of the *rho* gene, we constructed a scarless reporter system, in which the luciferase gene *luc* of firefly *Photinus pyralis* was fused to the *rho* promoter (P*_rho_*) at the position of the *rho* start codon. The P*_rho_-luc* fusion was similarly located in the *rho* locus of the chromosome in the wild type *B. subtilis* strain BSB1 and the *rho* deletion mutant RM (Bidnenko et al., 2017). The former strain (hereinafter WT) kept an active copy of the *rho* gene due to duplication of the *rho* locus during the construction. By comparing luciferase activity in WT and RM strains under different growth conditions, we sought to assess Rho autoregulation and possibly identify other regulatory factors.

In WT cells grown in rich medium LB, the expression of *rho* was mainly limited to the exponential growth phase, characterized by two peaks of luciferase activity at OD_600_ ∼0.15 and ∼0.4 (Fig 1A). In stationary phase, *rho* expression remained at a basal level. A low-level expression of *rho* and its down-regulation during the transition to stationary phase were reported previously (Ingham et al., 1999; Nicolas et al., 2012; Bidnenko et al., 2017; 2023). The inactivation of Rho in RM cells did not alter the kinetics of *rho* expression, but rather increased its level approximately threefold, suggesting a release from autoregulation (Fig 1A). WT cells grown in the sporulation-promoting Difco Sporulation Medium (DSM) showed a different *rho* expression kinetics, characterized by the additional peak of luciferase activity in stationary phase (Fig 1B). The reactivation of *rho* expression coincided in time with the induction of the sporulation-specific *spoIIAA-AB-sigF* operon, monitored in a separate strain using the P*_spoIIAA_-luc* transcriptional fusion (Fig 1B; Bidnenko et al, 2017). The similarity in timing suggested that *rho* expression during stationary phase in DSM might be linked to sporulation.

**Fig 1.**
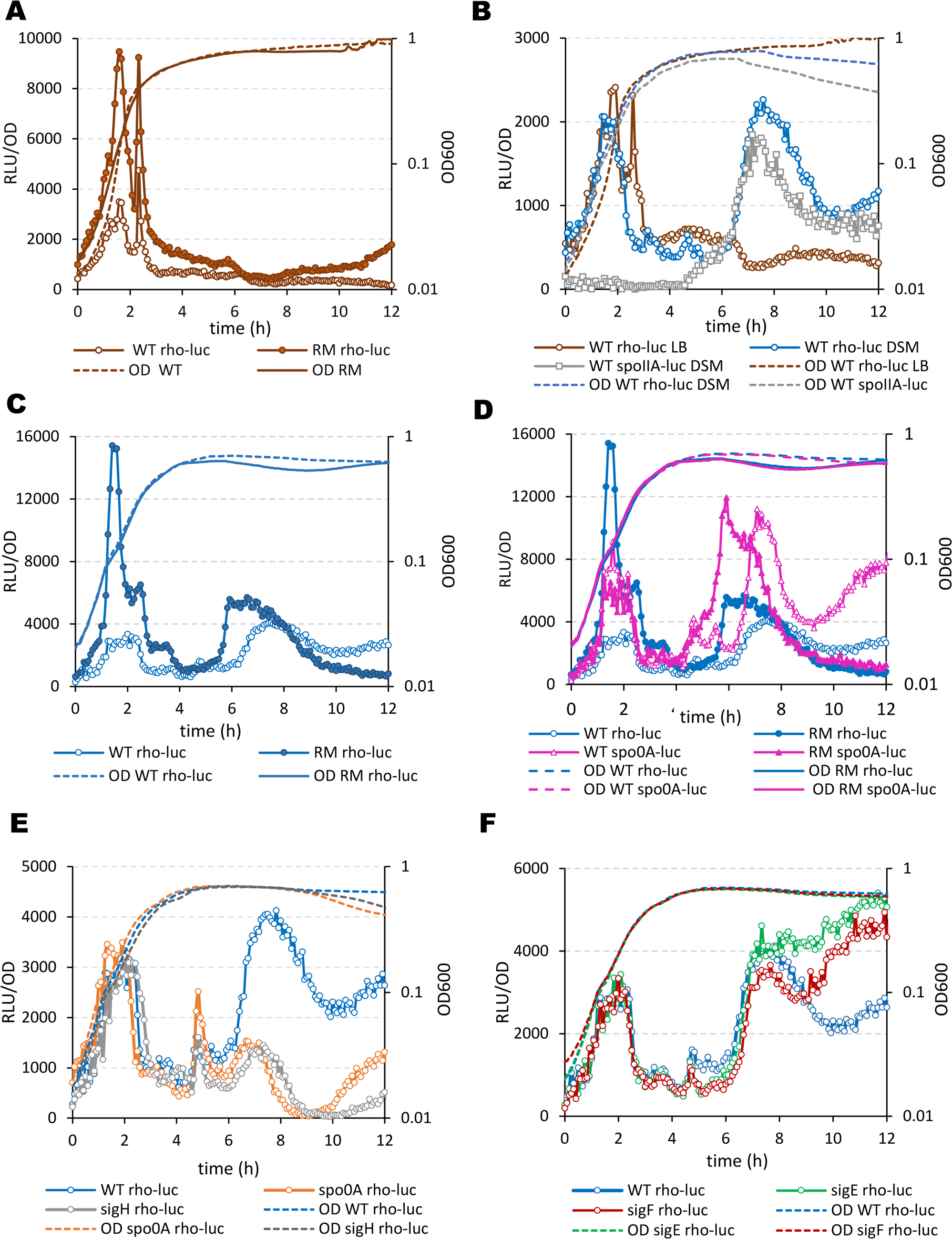
Rho is specifically expressed during *B. subtilis* sporulation. (A) Rho expression in vegetative cells is limited to the exponential growth phase and autoregulated. Kinetics of luciferase activity in *B. subtilis* WT P*_rho_-luc* (empty circles) and RM P*_rho_-luc* (filled-in circles) cells grown in rich medium LB. In this and other panels, the plain dotted and solid lines represent growth curves of the Rho-proficient strains and *rho*-deletion mutant (RM), respectively, measured by optical density OD_600_. (B) In the sporulation-inducing conditions, *rho* expression is additionally activated in stationary phase. Comparative analysis of the luciferase activity in WT P*_rho_-luc* cells grown in LB (brown circles) and in sporulation medium DSM (blue circles). Activation of luciferase expression from the early sporulation promoter *spoIIA* in the control WT P*_spoIIA_-luc* cells grown in DSM (grey squares) marks the initiation of sporulation. (C) In the sporulation-inducing conditions, the autoregulation of Rho is weakened in stationary phase. Kinetics of luciferase activity in *B. subtilis* WT P*_rho_-luc* (empty blue circles) and RM P*_rho_-luc* (filled-in blue circles) cells grown in sporulation medium DSM. (D) Rho expression in the sporulation-inducing conditions correlates with the activation of Spo0A. Comparative analysis of luciferase expression from P*_rho_-luc* (blue circles) and P*_spo0A_-luc* (pink triangles) transcriptional fusions in WT (empty symbols) and RM (filled-in symbols) cells grown in DSM. (E and F) Rho expression in stationary phase depends on the initiation of sporulation. Kinetics of luciferase expression from P*_rho_-luc* transcriptional fusions in WT cells (blue circles) and the sporulation mutants (*E*): *spo0A* (orange circles) and *sigH* (grey circles); and (*F*): *sigF* (red circles) and *sigE* (green circles). Measurements were taken every 5 minutes after cells inoculation in media at optical density OD_600_ ∼0.025 (time point 0). For each strain, plotted are the mean values of luminescence readings corrected for OD from four independent cultures analyzed simultaneously. Each panel presents the simultaneously collected data. The data in (C) and (D) and in (E) and (F) were obtained in two independent experiments. Each strain and growth condition was tested at least three times. The results from the representative experiment are presented.

In RM cells grown in DSM, luciferase activity increased ∼7-fold compared to WT cells during exponential growth, but only ∼1.5-fold in stationary phase. This suggests that *rho* expression in stationary phase is not subject to significant autoregulation. Curiously, however, the stationary phase-specific expression of *rho* was activated considerably earlier in RM than in WT cells (Fig 1C). This effect was reminiscent of the accelerated expression of *spo0A* gene in the sporulating RM cells caused by the up-regulation of Spo0A phosphorelay (Bidnenko et al., 2017). Therefore, we compared the expression of *rho* and *spo0A* genes in WT and RM cells grown in DSM using the P*_rho_*-*luc* and P*_spo0A_*-*luc* fusions (Mirouze et al., 2012; Bidnenko et al., 2017). In both strains, *rho* expression in stationary phase timely followed *spo0A* and, in RM cells, the maximal expression levels of both genes was reached ∼1.5 hour earlier than in WT (Fig 1D). This observation additionally argued for the sporulation-specific expression of *rho* as a function of Spo0A∼P activity.

To get more insights into the sporulation-specific expression of *rho*, we analyzed the P*_rho_*-*luc* activity in *B. subtilis* sporulation mutants. First, we considered *spo0A* and *sigH* mutations, which block initiation of sporulation by inactivating Spo0A and the transition phase-specific sigma factor SigH, respectively. Neither *spo0A* nor *sigH* mutations affected P*_rho_-luc* activity during exponential growth in DSM, but both specifically blocked it in stationary phase, thus confirming that *rho* expression at this stage is dependent on sporulation-specific factors (Fig 1E).

Next, we assessed the involvement of the alternative SigF and SigE factors controlling early gene expression in the forespore and mother cell, respectively. The kinetics of P*_rho_*-*luc* activity in *sigF* and *sigE* mutants was rather similar to WT cells. However, the stationary phase-specific luminescence appeared slightly lower in the *sigF* background and, in both mutants, remained high after reaching a maximal level (Fig 1F). These results indicated some deregulating effects of *sigF* and *sigE* mutations and suggested that *rho* isexpressed in both compartments of sporulating cells.

Considering that the known SigA-controlled P*_rho_* promoter is mainly active during exponential growth (Fig 1A), we sought for additional regulatory factors of *rho* expression during sporulation.

### The read-through transcription from the upstream SigH-controlled promoter contributes to *rho* expression during sporulation

The *rho* gene is transcribed from the cognate P*_rho_* promoter together with the upstream non-coding S1436 element and the downstream *rpmE* gene in a transcript of ∼1.9 kb (Quirk et al., 1991; Ingham et al, 1999; Nicolas et al., 2012). Additionally, the genome-wide transcriptome analyses have revealed several longer read-through transcripts at the *rho* locus, which initiate at the upstream *spo0F* and *fbaA-ywjH* promoters and bypass the intrinsic terminators of the *ywjH* and *glpX* genes (Nicolas et al., 2012; Mandell et al., 2022b; Fig 2A).

**Fig 2.**
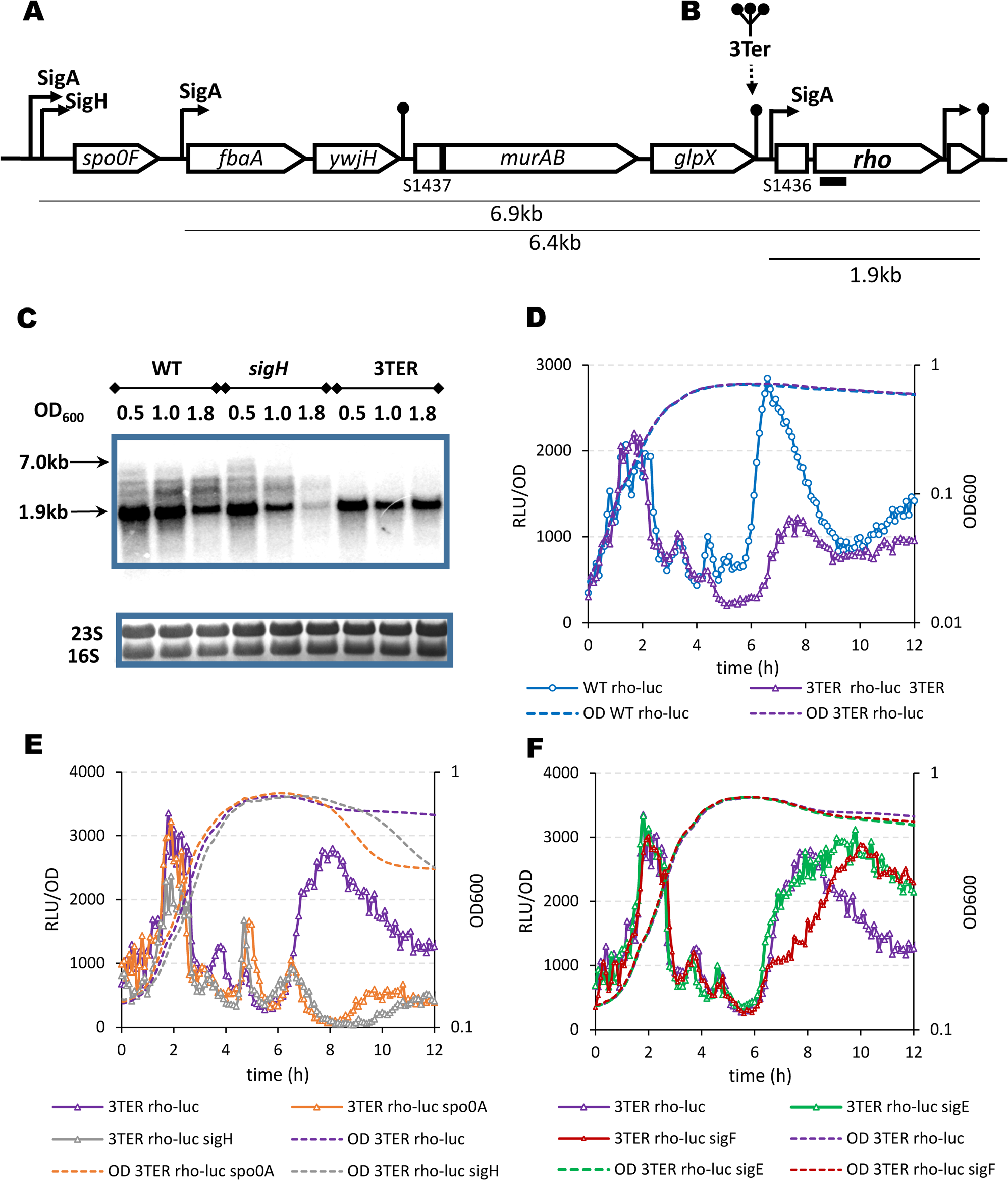
Role of the read-through transcription in the *rho* expression during sporulation. (A) Schematic representation of gene organization and transcription within the *rho* locus of *B. subtilis* chromosome. Arrow-shaped and flat rectangles indicate protein-coding genes and non-coding RNA S-segments (Nicolas et al., 2012), respectively. Curved arrow-ended lines represent promoters; their regulatory sigma factors are indicated. Straight dot-ended lines represent intrinsic terminators. The underlying lines indicate the transcripts of the *rho* gene, which were identified in (Nicolas *et al*., 2012); the transcript initiated at the known *rho* promoter is bolded. The approximate size of the transcripts is shown. Small black rectangle schematizes the 5’ *rho*-specific riboprobe used in the Northern blot. (B) Cartoon of the insertion of three intrinsic terminators at the end of *glpX* gene in the 3TER strain. (C) Northern blot analysis of the *rho*-specific transcripts in the wild type (WT), and the mutant *sigH* and 3TER strains during sporulation. Cells were grown in DSM and sampled during exponential growth (OD_600_ 0.5), the transition to stationary phase (OD_600_ 1.0) and in stationary phase (OD_600_ 1.8). Total RNA was extracted, processed and hybridized with the *rho*-specific riboprobe as described in Materials and Methods. *Upper panel*. The *rho*-specific transcripts visualized by Northern blotting. The arrows indicate the approximate size range of the transcripts. *Bottom panel*. Ribosomal 16S and 25S RNAs stained with ethidium bromide were used to control the equilibrium of the loaded RNA samples. (D) Suppression of the read-through transcription inhibits *rho* expression in stationary phase. Kinetics of luciferase activity in *B. subtilis* WT P*_rho_-luc* (blue circles) and 3TER P*_rho_-luc* (violet triangles) cells grown in the sporulation-inducing DSM. (E and F) In the absence of read-through transcription, the residual expression of *rho* in stationary phase still depends on sporulation. Kinetics of luciferase expression in 3TER P*_rho_-luc* strain (violet triangles) and its sporulation mutants (*E*): *spo0A* (orange triangles) and *sigH* (grey triangles); and (*F*): *sigF* (red triangles) and *sigE* (green triangles) grown in DSM. The data in (E) and (F) were obtained in the same experiment and are independent from (D). The data were collected and processed as described in Fig 1. The experiments were reproduced at least three times. The results from the representative experiment are presented.

We asked whether read-through transcription could play a role in the expression of *rho*, in particular during sporulation, as *spo0F* gene is known to be transcribed from alternative SigA- and SigH-dependent promoters, with the latter also being regulated by Spo0A∼P (Lewandoski et al., 1986; Predich et al., 1992; Strauch et al., 1993; Asayama et al., 1995). To this end, we sought to enhance transcription termination upstream of the *rho* gene and inserted a DNA fragment containing three intrinsic transcription terminators after the stop codon of the *glpX* gene (hereafter referred 3Ter; Fig 2B; Vagner et al., 1998; Bidnenko et al., 2017). To assess read-through transcription at the *rho* locus and the termination efficiency of the 3Ter insertion, we analyzed *rho*-specific transcripts in WT, *sigH* and 3TER mutant cells by Northern blot (Fig 2C). Cells were grown in DSM and sampled during exponential growth (OD_600_ 0.5), transition phase (OD_600_ 1.0) and in stationary phase, at OD_600_ 1.8. In WT and *sigH* mutant cells, we observed several *rho*-specific RNA species, the smallest and most abundant of which corresponded in size to the 1.9 kb *rho-rpmE* mRNA, and the largest (∼7 kb) to transcripts initiated at the *fbaA* and/or *spo0F* promoters. Other RNAs of ∼ 4-5 kb in size cannot be assigned to any defined promoter and may result from ribonuclease processing of the larger transcripts. During transition to stationary phase, the larger *rho*-specific RNAs accumulated in WT cells, but gradually disappeared in *sigH* mutant, in accordance with the notion that *spo0F* transcription in stationary phase depends on SigH activity (Fig 2C). Notably, none of the larger *rho*-specific transcripts were detectable in 3TER cells (Fig 2C) confirming that they do indeed originate from upstream of the inserted terminators. All together, these results show an efficient read-through transcription of the *rho* gene, mainly initiated at the *spo0F* promoter.

Northern analysis also provided information about P*_rho_* functioning. The levels of the 1.9 kb *rho* transcript decreased in stationary WT cells, in line with the down-regulation of the SigA-dependent P*_rho_* at this stage (Fig 2C; Nicolas et al., 2012), and became barely detectable in the stationary *sigH* mutant cells (Fig 2C). However, the dramatic decrease in the abundance of this short transcript in the *sigH* mutant was not due to the inhibition of read-through transcription from P*_spo0F_* promoter, as the amount of the 1.9 kb *rh*o transcript remained high in the stationary 3TER cells (Fig 2C).

To better understand the expression of *rho* in the absence of read-through transcription, we analyzed the activity of P*_rho_*-*luc* fusion in the 3TER strain and its sporulation-deficient derivatives. In 3TER P*_rho_*-*luc* cells grown in DSM, luciferase activity was identical to WT P*_rho_*-*luc* during exponential growth, but decreased ∼3-fold in stationary phase, highlighting the role of read-through transcription in the *rho* expression at this stage (Fig 2D). Intriguingly, the residual luminescence in the stationary 3TER P*_rho_*-*luc* cells was blocked by *spo0A* and *sigH* (Fig 2E) and partially inhibited by *sigF* inactivation, while mutation of *sigE* had no effect (Fig 2F). All together, these results indicated that the *glpX-rho* intergenic region contains additional structural factor(s) of *rho* expression active during sporulation and likely dependent on SigF.

### The sporulation-specific expression of *rho* is compartmentalized

Since SigF is primarily associated with the forespore, this led us to investigate potential compartmentalization of *rho* expression during sporulation. To this end, we constructed a P*_rho_*-*gfp* fusion expressing green fluorescent protein GFP at the *rho* chromosomal locus of WT cells and analyzed its activity during sporulation at a single-cell level by fluorescence microscopy. We observed an increase of fluorescence intensity in the forespores of the sporulating WT P*_rho_*-*gfp* cells immediately after asymmetric division (S1 Fig and Fig 3). This observation further suggested a particular role of SigF in the sporulation-specific expression of *rho*. Following it, we analyzed the activity of P*_rho_*-*gfp* fusion in *sigF* and *sigE* mutant cells.

**Fig 3.**
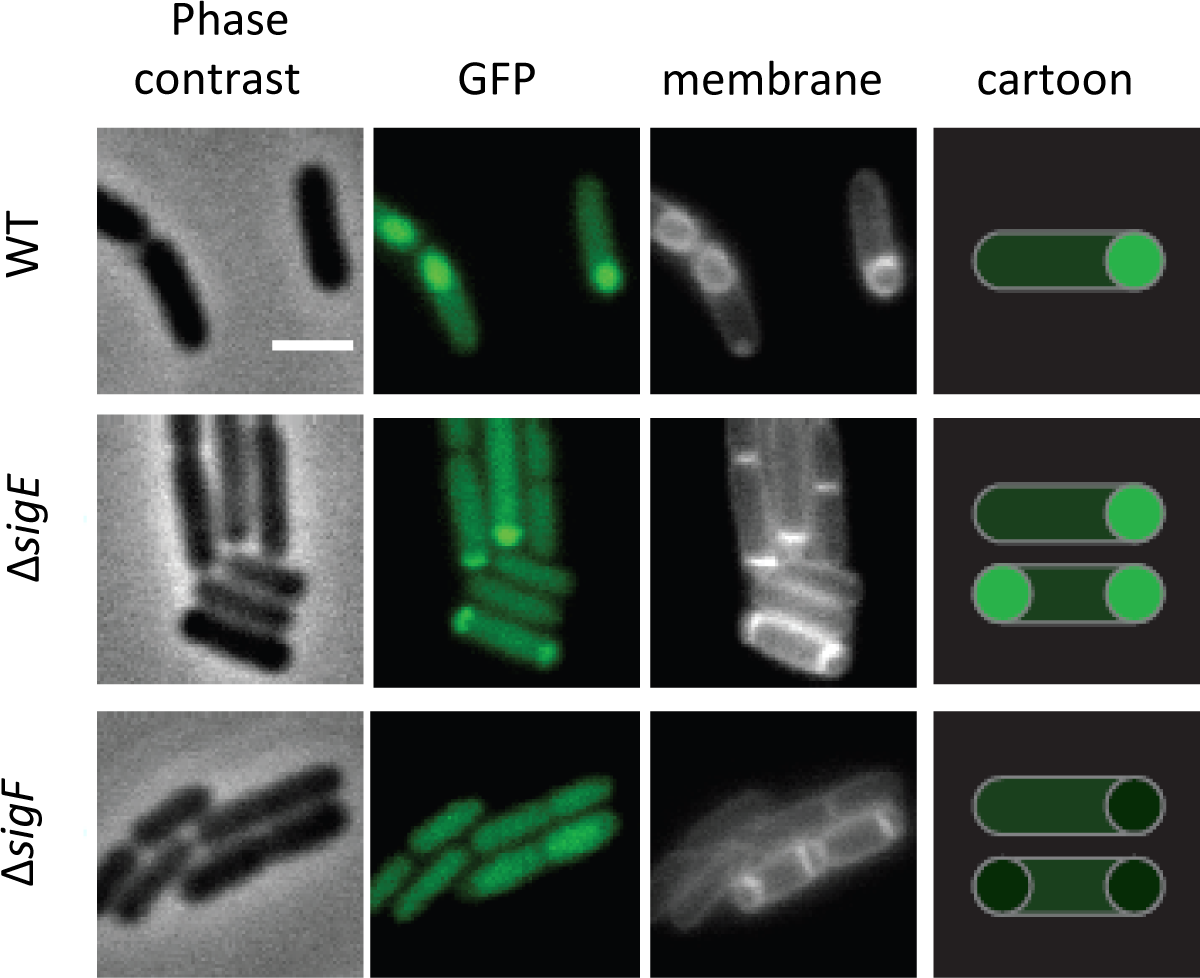
The expression of *rho* in the forespore compartment of sporangium depends on Sigma F. The WT P*_rho_*-*gfp* strain and its Δ*sigF* and Δ*sigE* mutant derivatives were induced for sporulation by the resuspension method as described in Materials and Methods. Cells were sampled three hours after resuspension in the nutrient-poor SM medium and observed by phase contrast microscopy, and by epifluorescence illumination of GFP or membrane-affine dye in two independent replicas. The right-hand cartoon depicts spatial GFP-mediated fluorescence in cells with asymmetric septa.

Absence of SigE disables the SigE-dependent inhibitors of septation, therefore provoking the formation of asymmetric septa at both poles of sporulating cells (Illing and Errington, 1991; Setlow et al., 1991; Lewis et al., 1994; Eichenberger et al., 2001). Because activation of SigE in the mother cell sequentially depends on SigF activity in the forespore (Londono-Vallejo & Stragier, 1995; and references therein), both *sigE* and *sigF* mutants display a similar disporic cell phenotype. Such disporic cells were readily seen in P*_rho_*-*gfp sigF* and P*_rho_*-*gfp sigE* sporulating cultures, in addition to cells containing one asymmetric septum. The mother cells of both *sigF* and *sigE* sporangia maintained GFP signal at a level similar to that of WT, indicating that *rho* expression is SigE-independent (Fig 3). In contrast, while the forespore-like structures showed an increased fluorescence in *sigE* mutant similarly to the WT forespores, their brightness was significantly diminished in *sigF* sporangia (Fig 3). Together with the results from the previous section, this observation indicated that, in forespores, *rho* is expressed from a promoter located in the *glpX-rho* intergenic region and dependent, directly or indirectly, on SigF.

### The 5’-UTR of *rho* contains a genuine SigF-dependent promoter active in forespores

Previous analyses have identified a *rho* promoter that matches the SigA-binding sequence consensus (^sigA^P*_rho_*) 293-320 bp upstream the *rho* start codon (Quirk et al., 1993; Ingham et al., 1999; Fig 4A). Recent reassessment of *B. subtilis* promoters using the unsupervised sequence clustering algorithm has classified ^sigA^P*_rho_* to the M16 cluster of SigA-dependent promoters, which are characterized by an extended 3’-terminal G-rich −10 box and a variable −35 box with the conserved TTG stretch (Nicolas et al., 2012; Fig 4B). Looking for alternative *rho* expression signals, we analyzed the *glpX-rho* intergenic region for the presence of the SigF-binding consensus −35 (GYATA) and −10 (GGnnAnAHTR) sequences, where Y is C or T; H is A or C or T; R is A or G; and n is any nucleotide (Wang et al., 2006). Such features were found within the ^sigA^P*_rho_* sequence itself (Fig 4C). Within the −10 box of ^sigA^P*_rho_*, we identified GGTAAAAATA sequence perfectly matching the SigF −10 consensus and, 14 bp upstream, a 5-nucleotide GAATA sequence differing from the SigF −35 consensus by one nucleotide. Of note, the same −35 sequence is present in the SigF-regulated promoters of *yuiC*, *yabT*, *yjbA*, *ypfB* and *ythC* genes (Wang et al., 2006). Moreover, the 14-bp spacer between −35 and −10 sequences is highly A/T-rich that is characteristic of most SigF-dependent promoters (Wang et al., 2006).

**Fig 4.**
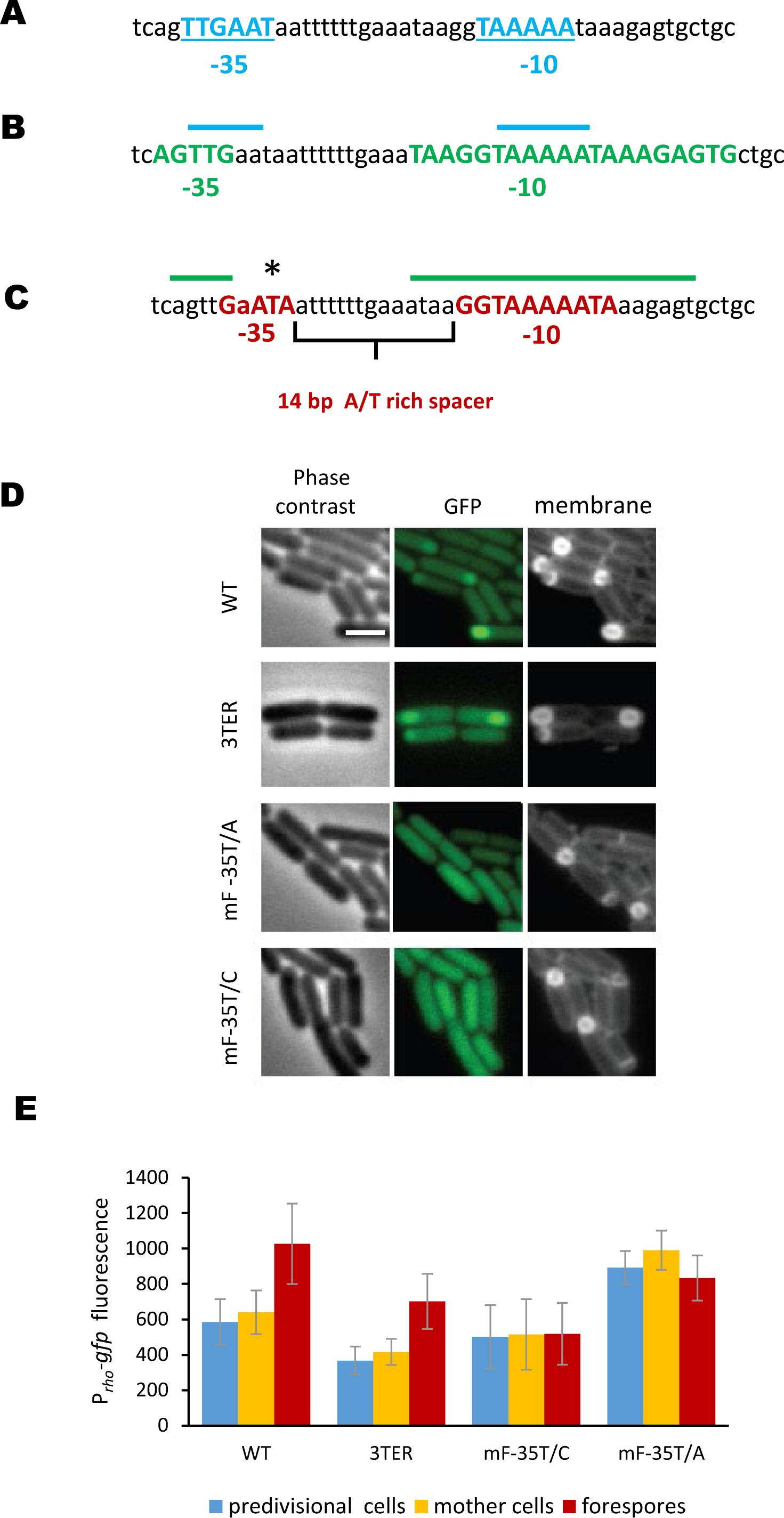
Forespore-specific expression of *rho* depends on the activity of a genuine SigF-dependent promoter. (A and B) Sequence and structural elements of the SigA-dependent *rho* promoter reported by: (A; Ingham et al., 1999) and (B; Nicolas et al., 2012). The −35 and −10 boxes are bolded and colored in bleu (A) and green (B). In (B), the upper blue lines indicate the relative position of the *rho* promoter identified by (Ingham et al., 1999). (C) Sequence and structural elements (bolded and colored in red) of putative SigF-dependent *rho* promoter identified in this analysis as matching consensus sequences recognized by SigF-containing RNA polymerase (Wang et al., 2006). The upper green lines indicate the relative position of the SigA-dependent *rho* promoter identified by (Nicolas et al., 2012). The asterisk indicates a conserved thymine nucleotide in the −35 sequence of the SigF-dependent promoter subjected to mutagenesis. (D and E) Mutations of the SigF-dependent *rho* promoter specifically inhibit *rho* expression in the forespores. The wild type (WT) and 3TER cells bearing the non-modified P*_rho_*-*gfp* fusion and WT P*_rho_*-*gfp* cells containing the indicated single-nucleotide mutations of putative SigF-dependent *rho* promoter were induced for sporulation and analyzed for GFP-mediated fluorescence as described in Materials and Methods and Fig 3. (D) Micrographs of typical cells observed by phase contrast, epifluorescence illumination of GFP or membrane-affine dye. (E) Average fluorescence intensities determined in predivisional cells and each compartments of sporangia in each strain. Fluorescence intensity was determined as described in Materials and Methods in two independent experiments. Bars represent standard deviation from the mean values.

To prove that the identified sequences constitute a novel SigF-dependent *rho* promoter (hereafter ^sigF^P*_rho_*), we proceeded to site-directed mutagenesis of the P_rho_-*gfp* fusion, anticipating that altering the SigF-binding sequences would specifically affect GFP expression in the forespores. Because of a strong overlap between the −10 sequences of ^sigA^P*_rho_* and putative ^sigF^P*_rho_*, we targeted the −35 GAATA sequence of the latter and replaced the invariant T nucleotide (Amaya et al., 2001; Wang et al., 2006) with A (*mF-35T/A* P_rho_-*gfp*) or C (*mF-35T/C* P_rho_-*gfp*) (Fig 4C). Next, we compared the activity of the mutant fusions with the original one (WT P_rho_-*gfp*) and the one preceded by the transcriptional terminators (3TER P_rho_-*gfp*). Cells were set to sporulate in DSM medium, and the fluorescence levels in predivisional cells and two compartments of sporangia were quantified at the time of a maximal SigF activity. As we observed earlier (Fig 3), the sporulating WT P_rho_-*gfp* cells showed a higher level of fluorescence in the forespores than in the mother cells; in the latter, *gfp* expression appeared similar to predivisional cells confirming our hypothesis that *rho* expression in the mother cell is SigE-independent (Fig 4D and 4E). A similar pattern of the brighter forespores was observed in 3TER P_rho_-*gfp* cells, even though the fluorescence was globally lower probably due to the inhibition of the read-through transcription by the 3Ter insert (Fig 4D and 4E).

In contrast, and as expected, both *mF-35T/A* and *mF-35T/C* mutations of P_rho_-*gfp* specifically inhibited the burst of fluorescence in the forespores, similarly to *sigF* mutation (Fig 4D and 4E, and Fig 3). At present, it is unclear why *mF-35T/A* P*_rho_*-*gfp* mutant strain displayed a higher basal fluorescence (Fig 4E). Nevertheless, these results clearly indicate that the forespore-specific expression of *rho* depends on the alternative promoter ^sigF^P*_rho_*.

This conclusion was further supported by the analysis of the mutated P*_rho_-luc* fusion: both point mutations *mF-35T/A* and *mF-35T/C* efficiently inhibited the sporulation-specific luciferase activity in 3TER cells (S2 Fig).

Overall, our results indicate that, during growth and sporulation, the expression of *rho* gene relies on three distinct features: (i) a SigA-dependent *rho* promoter that is active during exponential growth, (ii) read-through transcription from the upstream SigH-dependent *spo0F* promoter during early stationary phase and in the mother cells of sporangia, and (iii) a SigF-dependent *rho* promoter active in the forespores.

### Differential expression of Rho affects spore properties and morphology

The revealed spatiotemporal expression of *rho* suggested that besides modulation of Spo0A activity at the onset of sporulation (Bidnenko et al., 2017), Rho can be involved in the subsequent steps of spore development. To address this hypothesis, we used a set of five strains differentially expressing Rho during sporulation, namely: the wild type (WT), a strain blocked for the *rho* read-through transcription (3TER), their derivatives inactivated for *rho* expression in the forespore by *mF-35T/A* mutation of ^sigF^P*_rho_* (WT-mT/A and 3TER-mT/A), and a mutant deleted for *rho* (RM).

Initially, we compared sporulation capacities of the five strains. To this end, we induced sporulation by the resuspension method and assessed asymmetric division of individual cells and formation of the heat-resistant spores at the initial and final stages of sporulation, respectively. Among the five strains, RM cells showed a highest rate of asymmetric division corroborating our previous data on the accelerated sporulation of Δ*rho* mutant, most probably due to a more efficient activation of Spo0A (S3 Fig; Bidnenko et al., 2017). Interestingly, the proportion of cells with asymmetric septum was also higher in 3TER and 3TER-mT/A strains compared to WT or WT-mT/A mutant (S3 Fig) suggesting a regulatory crosstalk between Spo0A∼P activating read-through transcription of *rho* and Rho modulating activity of the Spo0A phosphorelay. However, while RM formed the heat-resistant spores considerably faster and with maximal yield, other mutants resembled more WT cells (S3 Fig). Therefore, only the complete inactivation of Rho significantly affected sporulation dynamics.

Next, we let cells to sporulate in DSM for 24 hours, during which more than 80 percent of cells in each strain formed spores, purified the mature spores, and compared some of their damage resistance properties. It should be noted, however, that most resistance phenotypes of spores are determined by multifactorial mechanisms, what makes their dissection difficult (reviewed in Setlow, 2006; Setlow and Christie, 2023). All spores differentially expressing *rho* were similarly resistant to lysozyme, which targets the cortex peptidoglycan (reviewed in Henriques and Moran, 2007), and became similarly sensitive to it after chemical removal of the coat (S4 Fig). This indicates that the permeability of spore coat to lysozyme was not detectably affected by altered expression of *rho*. Likewise, all five types of spores showed similar levels of wet-heat resistance, a complex spore phenotype determined by factors expressed in both spore compartments (S4 Fig). At the same time, unlike the others, RM spores contained higher (∼35%) levels of dipicolinic acid (DPA) (Fig 5A), an important factor of the heat-resistance synthesized in the mother cell, but accumulated in the spore core (reviewed in Nicholson et al., 2000; Setlow, 2016). Moreover, the mutant RM and also WT-mT/A and 3TER-mT/A spores, formed in the absence of the forespore-specific expression of *rho*, appeared more sensitive to ultraviolet (UV) light than WT and 3TER spores (Fig 5B). This effect was spore-specific, as vegetative RM and WT cells were equally resistant to UV radiation (S5 Fig). The UV-resistance of mature spores is mainly determined by the α/β small acid-soluble proteins (SASPs) and photoproduct lyase SplB synthesized in the forespores (Mason and Setlow, 1986; Pedraza-Reyes et al., 1994; Setlow, 2007). Based on our results, it is possible that Rho expressed from SigF-dependent promoter is involved in the regulation of some of these UV-resistance factors. Finally, we analyzed the ultra-structure of the purified spores by transmission electron microscopy, which revealed significant differences between the spores in the structure of their coats (Fig 6). In accordance with numerous previous studies (reviewed in Henriques & Moran, 2007; McKenney et al., 2013; Driks and Eichenberger, 2016), WT spores exhibited a regular surface layer composed of a lamellar inner coat and a striated thick electron-dense outer coat. Two coats were tightly attached to each other in a large majority (over 92 percent) of WT spores or occasionally separated across a limited area (Fig 6). In contrast, RM spores contained a lamellar inner coat similar to WT spores, but their outer coat remarkably lost electron-density and striated structure and appeared half thinner compared to WT spores (respectively, 36.2 +/− 0.0091 nm and 74.2 +/− 0.021 nm, as measured for 100 spores of each strain). Moreover, above 80 percent of RM spores showed an extensive detachment of inner and outer coats, often along the entire spore perimeter (Fig 6). Spores differentially expressing *rho* exhibited intermediate ranges of structural alterations of their coats. The outer coat of 3TER spores appeared less striated and low electron-dense compared to WT, therefore resembling the RM coat. Moreover, the outermost layer often peeled off the 3TER spore surface. However, we did not observe 3TER spores with the separated inner and outer coats (Fig 6). In WT-mT/A spores, the outer coat was electron-dense and striated like a WT layer, but detached from the inner coat in ∼30 percent of spores, although across less extended areas than in RM spores (Fig 6). Finally, 3TER-mT/A spores contained a low structured outer coat locally separated from the inner coat in ∼30 percent of spores (Fig 6). By combining these features, 3TER-mT/A spores appear most similar to RM spores, which seems in line with the absence of sporulation-specific *rho* expression in both mutant strains. We conclude that altering expression of *rho* during sporulation in the mother cell (3TER) or in the forespore (mT/A) induces specific defects of spore morphology, which are accentuated by the entire loss of the Rho activity in RM strain.

**Fig 5.**
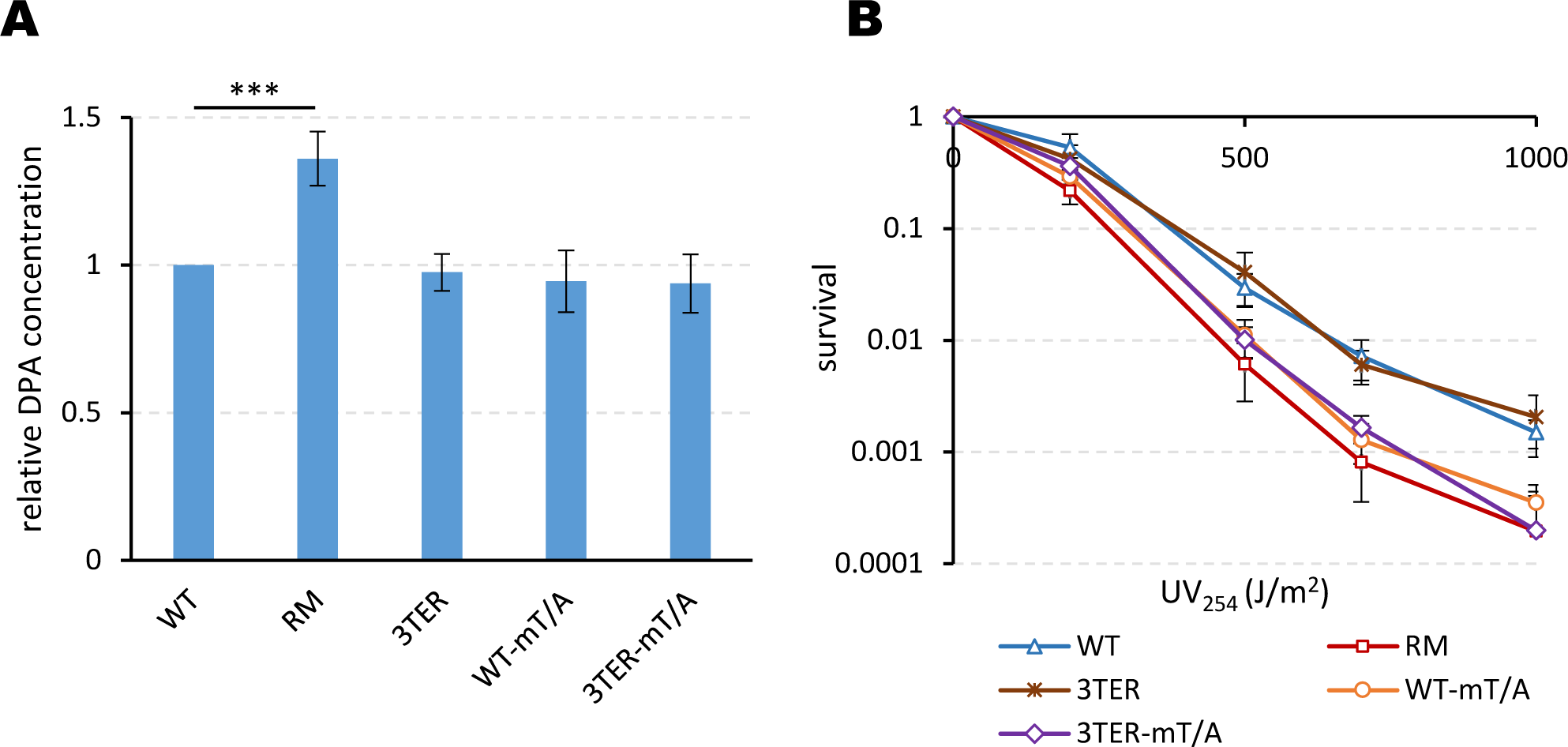
The alteration of the spatiotemporal expression of *rho* affects the resistance properties of mature spores. Spores produced by WT and 3TER cells expressing *rho* from the non-modified promoters, their mutant derivatives WT-mT/A and 3TER-mT/A inactivated for *rho* expression in the forespore and RM cells expressing no Rho were analyzed for the levels of dipicolinic acid (A) and the resistance to ultraviolet irradiation (B) as described in Materials and Methods. (A) Spore DPA contents are normalized to the wild type level. The assay was reproduced ten times with two independently prepared sets of five spores. Plotted are mean values from all measurements. Statistical significance was estimated with a two-tailed t-test. The displayed p-value is as follows: *** p ≤ 0.001. (B) UV test was reproduced five times with one set of spores. The bars represent standard deviation from the mean values.

**Fig 6.**
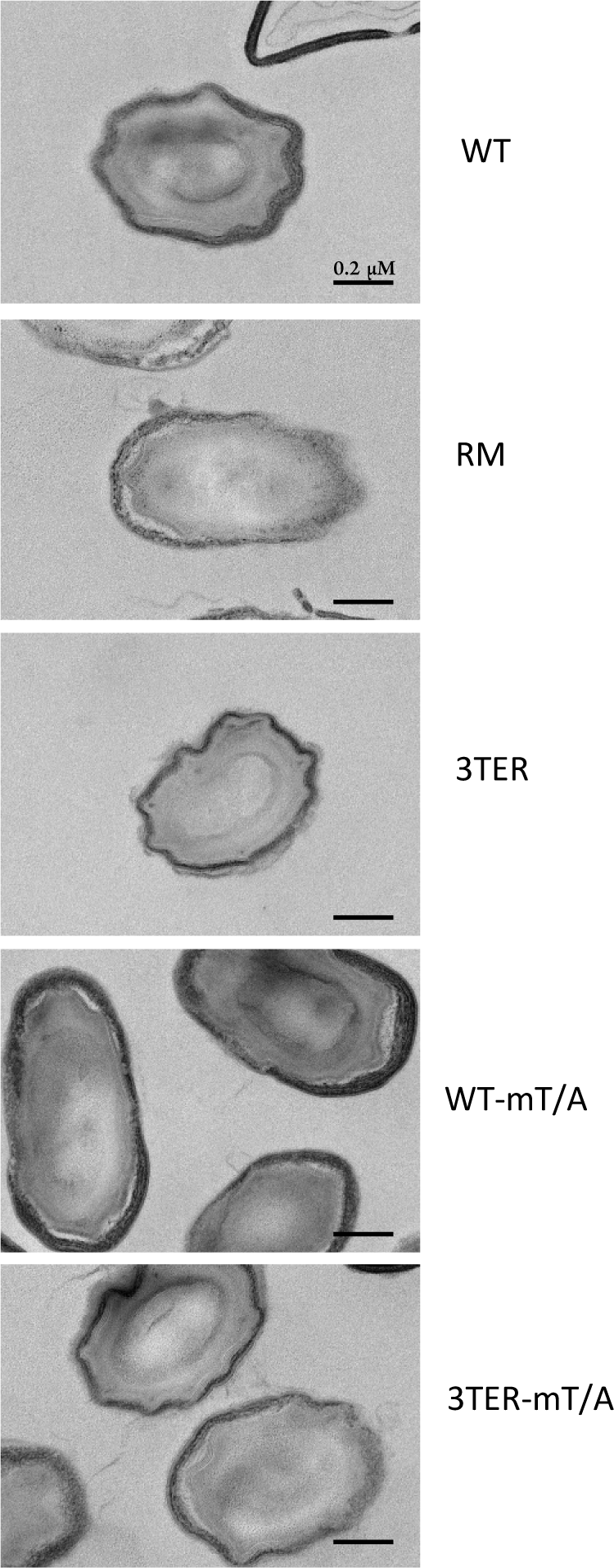
The alteration of the spatiotemporal expression of *rho* affects the morphology of spores. Thin section transmission electron micrographs of spores produced by cells differentially expressing *rho*. White arrows indicate a thinner and less electron-dense outer coat in RM, 3TER and 3TER-mT/A spores, which did not express *rho* in the forespores during maturation. Black arrows indicate the detachment of outer and inner coats in RM, WT-mT/A and 3TER-mT/A which did not express *rho* in the mother cells.

### Rho mediates spore germination and outgrowth

The ability to resume vegetative growth in favorable nutrient conditions, referred to as spore revival, is a basic spore property and spore quality indicator (Henriques and Moran, 2007; Setlow, 2013; Sinai et al, 2015; Ramírez-Guadiana et al., 2017a; Christie and Setlow, 2020). We compared the revival abilities of spores differentially expressing Rho by evaluating the optical density (OD_600_) of spore suspensions after the induction of germination. Rapid rehydration and structural changes of the germinating spores alter their optical properties resulting in a decrease in OD_600_. The OD_600_ of reviving spores remains stable during the post-germination phase of metabolic resumption and molecular reorganization, designated the “ripening period”, before increasing later during spore outgrowth, when the emerging cells start to grow and divide (Segev et al., 2013; Rosenberg et al., 2015). For simplicity, we consider the outgrowth period as a time needed for a spore culture to restore its initial OD_600_ after germination.

After the addition of the germinant L-alanine, RM spores lost OD_600_ faster than the others, thus showing the highest germination rate (Fig 7A). Similarly, RM spores germinated more efficiently with the germinants AGFK (a mixture of L-asparagine, glucose, fructose and potassium chloride) and L-valine recognized by other germinant receptors than L-alanine (S6 Fig). Other spores germinated similarly to WT with all three germinants (Fig 7A and S6 Fig). In contrast to the rapid germination, the outgrowth of RM spores in a defined minimal medium was significantly delayed compared to other spores (Fig 7B). At the same time, the viability of RM spores during germination was not decreased compared to WT spores (S7 Fig) and therefore could not explain the outgrowth delay. As for other spores, the enrichment of the growth medium by casamino acids (0.5%) increased the outgrowth rate of RM spores, which, nevertheless, remained below the WT level (Fig 7C). These results indicate that *rho* inactivation imposes stricter requirements on the spore outgrowth.

**Fig 7.**
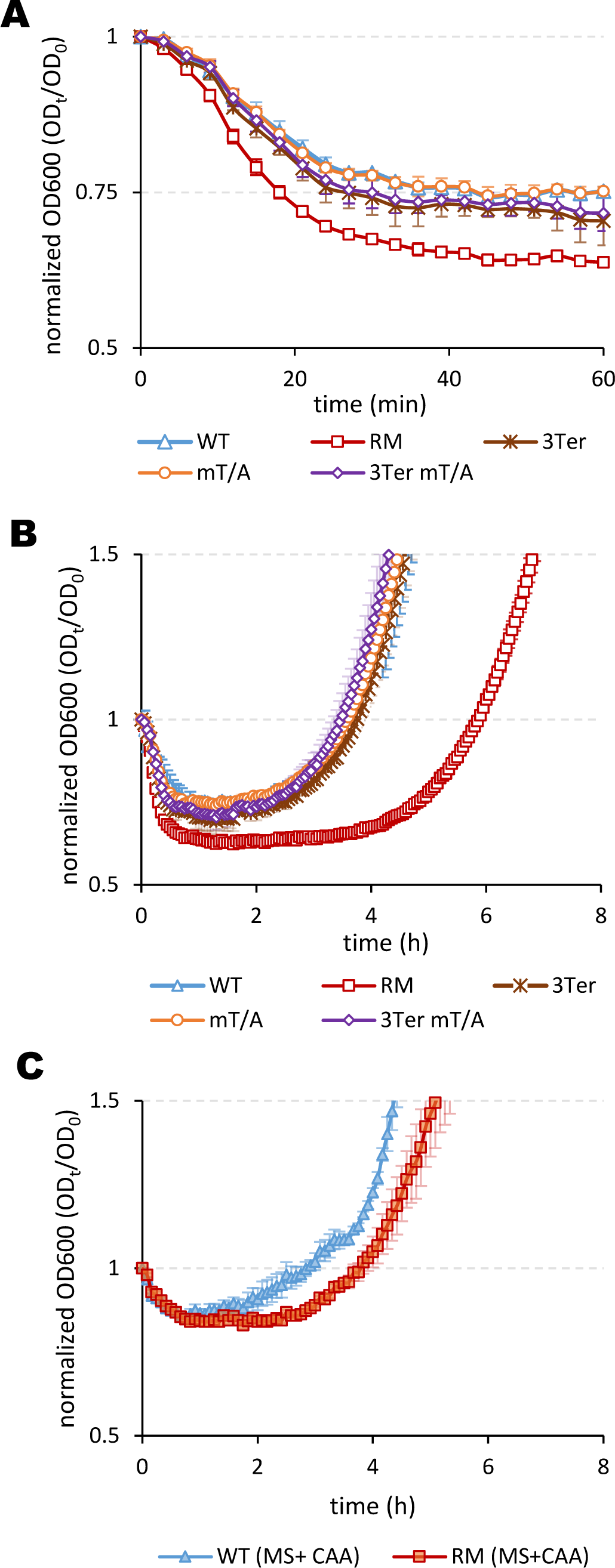
Rho activity determines the revival properties of spores. Spores produced by WT (blue triangles), RM (red squares), 3TER (brown crosses), WT-mT/A (orange circles) and 3TER-mT/A (violet diamonds) cells were induced by the germinant L-alanine (10mM) and analyzed for germination (A) and outgrowth in the nutrient-poor MS medium (B) as described in Materials and Methods. (C) WT (filled-in blue triangles) and RM (filled-in red squares) spores were compared for outgrowth in the nutrient-replenished MS medium containing 0.5% casamino acids. The experiments were performed at least twice with three independent sets of spores. Each experiment included up to six replicas of individual suspensions of spores. The results of the representative experiment are plotted. The bars represent standard deviation from the mean values.

We asked whether Rho activity is needed exclusively during spore morphogenesis for the formation of an appropriate molecular basis for spore revival in the future, or for a correct metabolic resumption during spore outgrowth. To answer this question, we placed the *rho* gene under the control of the IPTG-inducible promoter P_spac_, which allowed us to selectively control *rho* expression during sporulation and/or spore outgrowth. By immunoblot analysis of P_spac_-*rho* cells using the Rho^Bs^-specific antiserum, we established that the induction of P_spac_-*rho* by IPTG (100µM) determines a near-natural level of Rho protein (Materials and Methods; S8 Fig). Following this, we allowed P_spac_-*rho* cells to sporulate in the presence or absence of IPTG 100µM, purified the mature spores from the IPTG-induced and non-induced cultures and compared the revival of two types of P_spac_-*rho* spores in the conditions where *rho* expression was omitted or *de novo* induced by IPTG.

We observed that spores, which were formed in the absence of Rho (P_spac_-*rho* ^(-IPTG)^), exhibited an accelerated germination like RM spores, while those that expressed *rho* during morphogenesis (P_spac_-*rho* ^(+IPTG)^), germinated as WT (Fig 8A). Addition of IPTG together with the germinant did not modify germination kinetics of either spore type (Fig 8B). The two types of spores also behaved differently during outgrowth. In the absence of IPTG, the outgrowth of P_spac_-*rho* ^(-IPTG)^ spores was identical to RM spores, while P_spac_-*rho*^(+IPTG)^ spores grew out at an intermediate rate between WT and RM (Fig 8C). Interestingly, *de novo* induction of *rho* expression by IPTG had a small but reproducible stimulatory effect on the outgrowth rates, so that P_spac_-*rho*^(+IPTG)^ spores were able to “catch up” with the outgrowth of WT spores (Fig 8D). IPTG had no effect on the germination or outgrowth of WT and RM spores.

**Fig 8.**
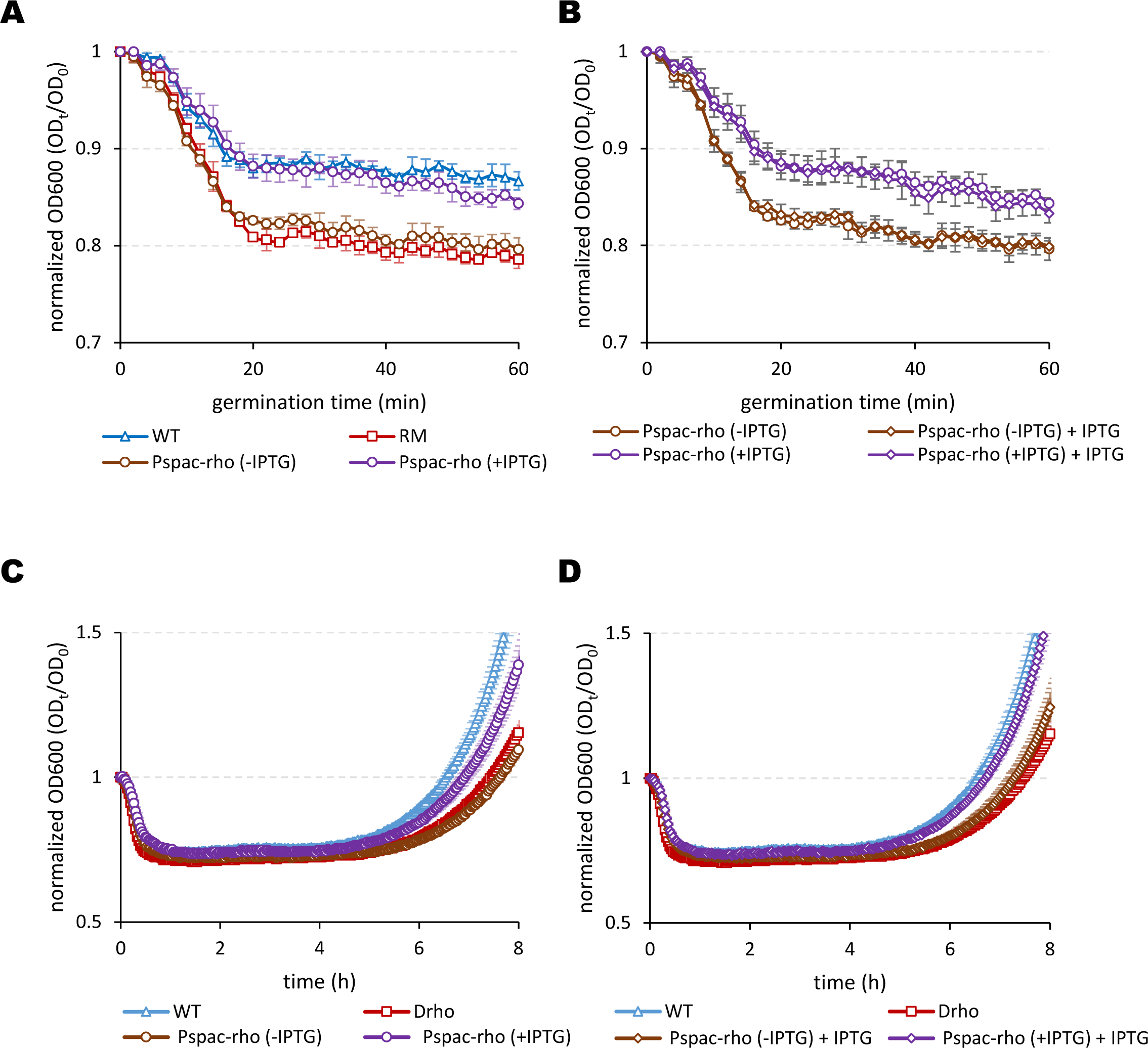
The wild-type rate of spore outgrowth requires the expression of *rho* during spore morphogenesis and *de novo* after germination. *B. subtilis* P*_spac_-rho* cells expressing Rho protein from the IPTG-inducible promoter were induced to sporulate in DSM in the absence (brown symbols) or presence (violet symbols) of IPTG 100µM. The mature spores were purified and analyzed for germination with L-alanine (A and B) and outgrowth in MS medium (C and D) in the absence (A, B and C; circles) or presence of IPTG 100mM (B and D; diamonds). The WT (bleu triangles) and RM (red squares) spores produced simultaneously with P*_spac_-rho* spores were used to compare the germination in (A) and outgrowth in (C and D). Of note, *rho* induction by IPTG (B): had no effect on the germination rates of the both types of P*_spac_-rho* spores, but (D): accelerated the outgrowth of spores expressing *rho* during morphogenesis (violet diamonds) to the WT level. The experiments were performed at least twice with two independent samples of the IPTG-induced or non-induced P*_spac_-rho* spores and included up to six replicas of individual suspensions of spores The results of the representative experiment are plotted. The bars represent standard deviation from the mean values.

From these experiments, we conclude that normal spore revival requires expression of *rho* during spore morphogenesis and, although to a lesser extent, *de novo* during spore outgrowth.

## Discussion

The main achievement of the present analysis is the revelation of spatiotemporal expression of the termination factor Rho during sporulation in *B. subtilis* via two distinct mechanisms: read-through transcription initiated at a distal promoter controlled by SigH in the mother cells, and transcription from the authentic SigF-dependent *rho* promoter in the forespores. These regulatory elements compensate for the inactivation of SigA-dependent *rho* expression at the end of exponential growth and allow the “refueling” of Rho protein in both compartments of the sporangium.

Our results also show that maintaining Rho levels during sporulation is necessary for the formation of spores with normal morphology, resistance properties and the ability to return to a vegetative state under favorable environmental conditions.

### Compartmentalized expression of *rho* during sporulation

The sporulation-specific read-through transcription of *rho* is initiated by the activation of the SigH-dependent promoter of *spo0F* gene encoding the phosphotransferase Spo0F of the Spo0A phosphorelay. The *spo0F* expression is regulated by Spo0A∼P and constitutes one of the positive feedback loops of Spo0A∼P at the onset of sporulation (Lewandoski et al., 1986; Predich et al., 1992; Strauch et al., 1993; Asayama et al., 1995; Chastanet et al., 2010). Previously, we have established that Rho negatively regulates the expression of some components of the Spo0A phosphorelay, like the sensor kinase KinB (Bidnenko et al., 2017). Therefore, correct modulation of Spo0A∼P activity at the initial steps of *B. subtilis* differentiation requires the decrease in the *rho* expression levels (Bidnenko et al., 2017). As we show here, this decrease is partially due to the silencing of the SigA-dependent *rho* promoter by yet unknown mechanism. It appears that alongside a self-reinforcing loop involving Spo0F, Spo0A∼P simultaneously activates the Rho-mediated regulatory circuit, which can be viewed as a negative feedback loop of Spo0A∼P. This regulatory function of Rho could be particularly important to fine-tune Spo0A∼P activity during initiation of sporulation and in the mother cells, where Spo0A∼P remains functional after asymmetric division (Fujita and Losick, 2003). Our data indicate that the mother cell-specific sigma factor SigE is not involved in the expression of *rho*. Nevertheless, inhibition of the *rho* read-through transcription results in the structural changes of spore coat, whose assembly depends on the activity of the SigE- and SigK-controlled genes (Eichenberger et al., 2003; Henriques and Moran, 2007; Driks and Eichenberger, 2016). This emphasizes an extended activity of Rho during the mother cell gene expression program. Identification of the alternative forespore-specific *rho* promoter matching the SigF-binding consensus allows us to classify *rho* as a member of the SigF regulon. Interestingly, an earlier analysis of forespore gene expression has detected the up-regulation of *rho* in sporulating *sigF*^+^ *B. subtilis* cells (see Supplemental Table1 in Wang et al., 2006). However, *rho* was not induced by the artificial expression of SigF during exponential growth in LB, a second criterion used for gene classification to the SigF regulon (Wang et al., 2006). It is possible that, in that analysis, the untimely activated SigF failed to compete with the abundant vegetative SigA for binding to the overlapping cognate *rho* promoters. The silencing of the SigA-dependent *rho* promoter might facilitate SigF binding. However, the mechanism of this silencing is unclear, and despite some discrepancies between the reports, SigA remains active in the forespores (Lord et al., 1999; Fujita, 2000; Li et Piggot, 2001; Marquis et al., 2008; Riley et al., 2018; 2021). It is also notable that the *rho* gene is located within a chromosomal cluster of SigF-dependent genes activated the first during the forespore formation (Steil et al., 2005; Wang et al., 2006), what pinpoints its early expression. At the same time, our data indicate the role of Rho in the resistance of spores to ultraviolet radiation, the main factors of which, the α/β-type small acid-soluble spore proteins (SASPs) and the spore photoproduct lyase SplB, are controlled by the late forespore-specific sigma factor SigG (Mason and Setlow, 1986; Pedraza-Reyes et al., 1994; Setlow, 2001). Therefore, like the read-through transcription in the mother cells, the SigF-mediated *rho* expression appears to ensure Rho activity during the entire forespore-specific gene expression program. It can also mediate the packaging of Rho protein in mature spores, where it has been recurrently identified by mass spectrometric analysis (Swarge et al., 2020; Tu et al, 2020).

### Rho involvement in the spatiotemporal regulation of spore development

Two spore phenotypes, a decreased UV-resistance and a structurally altered outer coat, resulting from the complete inactivation of *rho*, or following the inhibition of *rho* expression in the forespores and in the mother cells, respectively, highlight the importance of the compartmentalized Rho-mediated transcription termination for spore development. Our previous transcriptome analysis of RM cells in stationary phase revealed disordered and untimely expression of many sporulation genes via their extensive sense or antisense read-through transcription or, more rarely, unscheduled activation of promoters (Bidnenko et al., 2023). The relevant gene expression changes are compiled in S1 Table and can be visualized at http://genoscapist.migale.inrae.fr/seb_rho/ (Bidnenko et al., 2023; Dérozier et al., 2021). We posit that at least some of the sporulation genes abnormally expressed in the stationary RM cells are subject to Rho-mediated regulation during sporulation. At the same time, we are aware that most of potential sporulation-specific targets of Rho are not yet expressed in stationary phase. For instance, the stationary phase transcriptome of RM cells does not provide an obvious explanation for the decreased UV-resistance of spores differentially expressing *rho*. Indeed, the observed upregulation of several *ssp* genes in the stationary RM cells, in particular *sspC*, is contrary to what one could expect if spores become sensitive to UV. It is therefore likely that the effect of Rho in stationary phase and in sporulating cells is different for some genes.

The existing transcriptome data probably provide the most straightforward mechanistic hints on the involvement of Rho in the regulation of spore coat morphogenesis. The prominent coat defects in RM and 3TER-mT/A spores resemble those induced by the inactivation of morphogenetic proteins CotO, CotH or CotG, or auxiliary proteins of coat assembly, superoxide dismutase SodA and holin YwcE (McPherson et al., 2005; Zilhão et al., 1999; Isticato et al., 2020; Freitas et al., 2020; Henriques et al., 1998; Real et al., 2005; Henriques and Moran, 2007). All of the abovementioned mutant spores exhibit a disordered, low electron-dense and thinner outer coat frequently detached from the inner layer. The protein kinase CotH and its target protein CotG were recently identified as the main morphogenetic factors determining the regular structure of spore coat (Nguyen et al, 2016; Freitas et al., 2020; Isticato et al., 2020; Di Gregorio Barletta et al., 2022). It has been shown that phosphorylated CotG assembles in the coat, while its non-phosphorylated form sequesters other coat proteins in the mother cell cytoplasm (Di Gregorio Barletta et al., 2022).

Interestingly, the expression levels of *cotG* are strongly upregulated in RM cells, due to the transcription upshift at, or close to, the SigK-dependent *cotG* promoter and the read-through transcription initiated at the upstream *alsR* gene (Nicolas et al., 2012; Bidnenko et al., 2023; S1 Table). The same read-through transcript appears antisense to the oppositely oriented neighboring genes *cotB* and *cotH*. We suggest that Rho participates in the control of the transcription of the *cotB-cotH-cotG* gene cluster during sporulation and that inactivation of the *rho* expression in the mother cells modifies the CotG-CotH protein balance important for correct assembly of CotG, thus leading to spore coat defects. Work is in progress to test this hypothesis.

Alongside the shared phenotypes linked to the inhibition of spatiotemporal expression of *rho*, RM spores are specifically characterized by the increased levels of DPA content. The synthesis of DPA and its transfer to the core across the spore membranes involves the protein products of both mother cell- and forespore-specific genes (Daniel and Errington, 1993; Ramírez-Guadiana et al., 2017b; Gao et al., 2022). None of the genes involved in the DPA metabolism are subject to unscheduled activation in RM cells in stationary phase. However, we detected strong induction of a defined SigA-dependent antisense promoter located within the SigG-regulated *spoVAA-spoVAB-spoVAC-spoVAD-spoVAEB-spoVAEA-spoVAF*-*lysA* operon, whose products are essential for DPA uptake into the core and release during germination (Tovar-Rojo et al., 2002; Gao et al., 2022; Nicolas et al., 2012; Bidnenko et al, 2023). In the wild type *B. subtilis* cells, this antisense promoter is weakly activated under some stress conditions and late in sporulation (Nicolas et al., 2012). Remarkably, in RM cells, the generated ∼5 kb-long antisense transcript extends across *spoVAA, spoVAB*, *spoVAC* genes and the *dacF*-*spoIIAA-spoIIAB-sigF* operon. It remains to be established whether this antisense transcript affects the expression of the genes on the opposite strand during RM sporulation.

### Rho involvement in spore revival

RM spores also differ from their siblings by their impaired revival capacity, showing a rapid germination followed by slow outgrowth. The rate of germination in response to different nutrient stimuli is mainly determined by the levels of corresponding germinant receptors in the spore membrane (Cabrera-Martinez et al., 2003; Ghosh et al., 2012; Chen et al., 2014). The RM stationary phase transcriptome showed unscheduled up-regulation of the SigG-controlled *gerBA-gerBB-gerBC* operon encoding the AGFK germinant receptor GerB (Corfe et al., 1994) and the SigK-controlled *gerPA-gerPB-gerPC-gerPD-gerPE-gerPF* operon, whose products facilitate the passage of nutrient germinants across the spore coat (Behravan et al., 2000). Although these features suggest that Rho controls both operons during sporulation, they seem insufficient to explain the rapid germination of RM spores in response to L-alanine or L-valine. Among the *gerAA-gerAB-gerAC* genes encoding the L-alanine germinant receptor GerA (Feavers et al, 1990; Amon et al., 2022, and references therein), only *gerAA* appears up-regulated ∼3-fold in the stationary RM cells.

Interestingly, Rudner and colleagues identified *rho* among the genes, whose inactivation by Mariner transposon results in a premature germination of the developing spores triggered by the GerA receptor, probably in response to some undefined defects of spore envelope (Meeske et al., 2016; Ramírez-Guadiana et al., 2017). The authors report that GerA activity results in a decreased sporulation efficiency of the *rho::Mariner* mutant, which is opposite to our recurrent observation of a highly efficient sporulation in the *rho*-deleted RM cells. Nevertheless, the established genetic interaction between Rho and GerA proteins may be relevant to the germination phenotype of the mature RM spores, whose coat is severely altered. Moreover, the structure of RM spore coat suggests a lower level of coat protein cross-linking (Henriques and Moran, 2007), which can facilitate spore germination, as previously observed (Abhyankar et al., 2015). The germination rate of RM spores might be also influenced by high levels of DPA, a potent non-nutrient germinant that stimulates cortex hydrolysis by CwlJ hydrolase both directly and during nutrient-induced germination (Paidhungat and Setlow, 2000; Paidhungat et al., 2001).

More generally, spore germination is a complex highly heterogeneous process influenced by multiple factors, including, among others, the metabolic activity of cells during sporulation and the duration of spore formation (Segev et al. 2012; Bressuire-Isoard et al., 2018; Mutlu et al, 2018, 2020; Riley et al., 2021; Rao et al., 2022). From this viewpoint, a more synchronous and rapid sporulation in RM population (Bidnenko et al., 2017, and this study) might result in a more homogeneous molecular composition of spores and, consequently, in a decreased heterogeneity of spore germination.

At the same time, the absence of Rho negatively affects the outgrowth of the germinated RM spores even in a nutrient-enriched medium. Therefore, *rho* inactivation imposes more stringent requirements for spore outgrowth, which in itself is more exacting than vegetative growth (O’Brien and Campbell, 1957). Taken together, a rapid germination and a slow outgrowth of RM spores suggest that, in the wild type background, Rho determines the fitness of reviving spores.

Using a controlled *rho* expression system, we showed that the normal rate of spore outgrowth requires *rho* expression both during spore formation and after spore germination. During sporulation, Rho would ensure a proper composition of a spore molecular cargo required for spore revival, which also includes Rho protein itself (Mutlu et al, 2018; Rao et al., 2022; Swarge et al., 2020). After germination, Rho would be required in the ripening period, to control a proper termination of the actively resuming transcription, and later, during implementation of the transcription program of vegetative growth. This is consistent with the differential gene expression analysis of the reviving *B. subtilis* spores, which detected the upshifts in *rho* expression shortly after germination and at the end of outgrowth (Swarge et al., 2020). Most probably, the resumption of *rho* expression during outgrowth reflects the reactivation of SigA-dependent *rho* promoter.

To conclude, this and our previous analyses trace the activity of Rho through each stage of the complex program of *B. subtilis* sporulation, from the regulation of Spo0A∼P activation at the onset of sporulation, through the subsequent morphogenesis of spores and up to the modulation of spore revival. This highlights the existence of specific targets for Rho-mediated transcription termination within the regulon of each alternative Sigma factor, which certainly include, alongside the known coding genes, a particular set of non-coding and anti-sense transcripts. Appropriate regulation of these targets requires in turn an accurate time- and spatial-specific regulation of the *rho* expression, through mechanisms we describe. Overall, our results strengthen our vision of Rho as a built-in regulatory module of *B. subtilis* cell differentiation.

## Materials and Methods

### Bacterial strains and growth conditions

All strains used in the analysis originate from *B. subtilis* 168 *trp^+^* strain BSB1 (Nicolas et al., 2012) and are listed in S2 Table. The *Escherichia coli* TG1 strain was used for construction of intermediate plasmids. Cells were routinely grown in liquid or solidified rich LB medium at 37°C. Standard protocols were used for transformation of *E. coli* and *B. subtilis* competent cells (Harwood and Cutting, 1990). Sporulation of *B. subtilis* cells was induced by the method of nutrient exhaustion in supplemented Difco sporulation medium (DSM; Difco) (Schaeffer et al., 1965) or by the resuspension method (Harwood and Cutting, 1990), as detailed in (Bidnenko et al., 2023). Optical density of the bacterial cultures was measured with NovaspecII Visible Spectrophotometer, Pharmacia Biotech. When required, antibiotics were added at following concentrations: 0.5 *μ*g per ml of erythromycin, 3 µg per ml of phleomycin, 100 *μ*g per ml of spectinomycin, and 5 *μ*g per ml of chloramphenicol to *B. subtilis* cells; and 100 *μ*g per ml of ampicillin or 20 µg per ml of kanamycin to the plasmid-containing *E. coli* cells. For the induction of P*_spac_-rho* fusion, IPTG (isopropyl-β-D-1-thiogalactopyranoside) inducer was added to cells and spores to final concentration 100µM.

### Strains and plasmids construction

All intermediate plasmids were constructed using Q5 High Fidelity DNA Polymerase and DNA modification enzymes purchased from New England Biolabs. Transcriptional fusion of the luciferase gene *luc* and the *rho* promoter (P*_rho_-luc*) was constructed as follows. The oligonucleotide pairs glpXBam/veb738 were used to amplify a ∼1 Kb fragment of *B. subtilis* chromosome located directly upstream the *rho* start codon. The 5’ part of the *luc* gene was amplified from the plasmid pUC18cm-Luc (Mirouze et al., 2012) using oligonucleotides *lucintrev* and veb739 complementary to veb738. Two DNA fragments were joined by PCR using the primers glpXBam and *lucintrev*, and the resulting DNA fragment was cloned at pUC18cm-Luc using BamHI and BstBI endonucleases and T4 DNA ligase. The resulting plasmid pBRL862 was integrated by single crossover at the *rho* chromosomal locus of BSB1 (WT) and BRL1 (Δ*rho*) cells using selection for chloramphenicol-resistance. By this event, the *luc* gene was placed under all natural regulatory signals of the *rho* expression. In the WT background, the replaced intact copy of *rho* gene remained active under the control of own promoter.

Insertion of three intrinsic transcription terminators (3TER) upstream the chromosomal P*_rho_-luc* fusion was performed as follows. The 3TER DNA fragment amplified from pMutin4 plasmid using oligonucleotides veb734 and veb735 was end-joined with the DNA fragments amplified from pBRL862 using the pairs of oligonucleotides glpXBam/veb798 (5’-complementary to veb734) and *lucintrev*/veb797 (5’-complementary to veb735). The primers glpXBam and *lucintrev* were used in the joining reaction. The PCR product was cut by EcoRV and BstBI endonucleases and cloned at a similarly cut and gel-purified pBRL862 plasmid. The resulting plasmid pBRL1107 was integrated at the *rho* locus of BSB1 chromosome as above. The created 3TER-P_rho_*-luc* transcriptional fusion contains intact 5’-UTR of *rho* and the 3TER insertion immediately after the *glpX* stop codon.

Transcriptional fusion between the *gfp* gene, encoding Green Fluorescent Protein, and the *rho* promoter (P*_rho_-gfp*) was constructed similarly to P*_rho_-luc*. The oligonucleotides glpXBam and veb742 were used for amplification of the *rho* moiety from BSB1 chromosome, and the *gfp* gene was amplified from the plasmid pCVO119 using the primers veb741 and veb740 complementary to veb742. The primers glpXBam and veb741 were used for joining PCR, and the resulting DNA fragment was cloned in pCVO119 plasmid using BamHI and NcoI endonucleases and T4 DNA ligase. The resulting plasmid pBRL893 was integrated by single crossover at the *rho* locus of the BSB1 chromosome using selection for spectinomycin resistance.

To construct the 3TER-P*_rho_-gfp* transcriptional fusion, the DNA fragment containing 3TER and the rho 5’-UTR was amplified from pBRL1107 plasmid using the primers glpXBam and veb742 and fused to the *gfp* gene as described above for pBRL893. The product of joining PCR was cut by EcoRV and NcoI endonucleases and cloned at a similarly cut and gel-purified pBRL893. The resulting pBRL1150 plasmid was inserted in the BSB1 chromosome as above. Site-directed mutagenesis of the Sigma F-dependent *rho* promoter of the P*_rho_-luc* and P*_rho_-gfp* transcriptional fusions was performed as follows. The plasmids pBRL893 and pBRL1107 were entirely amplified using the pairs of side-by-side oligonucleotides veb803/veb802, containing a 5’-terminal C nucleotide to introduce the point mutation *mF-35T/C*, and veb803/veb805, with a 5’-terminal A for the point mutation *mF-35T/A*. The PCR products were phosphorylated using T4 polynucleotide kinase, self-ligated and treated with DpnI endonuclease to remove the template DNA prior transformation in *E. coli* cells. The resulting mutated derivatives of the plasmids pBRL893 (pBRL1141 and pBRL1164) and pBRL1107 (pBRL1116 and pBRL1162) were controlled by sequencing and inserted into the *B. subtilis* BSB1 chromosome as above to produce the mutated fusions *mF-35T/C* P*_rho_*-*gfp* and *mF-35T/A* P*_rho_*-*gfp* and 3TER *mF-35T/C* P*_rho_*-*luc* and 3TER *mF-35T/A* P*_rho_*-*luc*, respectively.

Structural modifications of the *rho* expression unit at natural locus were performed by the method for allelic replacement using a shuttle vector pMAD (Arnaud et al., 2004).

To insert 3TER transcription terminators at the *rh*o locus, two DNA fragments amplified from the BSB1 chromosome with the pairs of oligonucleotides veb795/veb796 and veb797/veb798 were end-joined to the 3TER DNA fragment (see above) by PCR using the primers veb795 and veb796. The product of joining PCR was cut with BamHI and EcoRI endonucleases and cloned in the pMAD vector (Arnaud et al., 2004). The resulting plasmid pBRL1108 was integrated into the *rho* locus of the BSB1 chromosome with selection of the erythromycin-resistant transformants at 37°C non-permissive for plasmid replication. Single transformants were propagated in LB without antibiotic at 30°C to induce the loss of the vector and platted at LB-plates at 37°C. The erythromycin-sensitive clones were selected among the grown colonies and tested by PCR for the presence of the 3TER insertion using oligonucleotides veb734 and veb735, and the selected clones were controlled for structural integrity of the *glpX-3TER-rho* region by sequencing.

The point mutation *mF-35T/A* was introduced in the *rho* locus of BSB1 and BRL1130 (3TER) strains as follows. Two DNA fragments were amplified from pBRL1164 plasmid and BSB1 chromosome by PCR with the respective pairs of oligonucleotides veb796/veb808 and veb806/veb807, complementary to veb808. The fragments were joined by PCR using veb796 and veb806 as primers, and the resulting product was cloned at pMAD vector using BamHI and EcoRI endonucleases and T4 DNA ligase. The resulting plasmid was transformed into BSB1 cells and the transformants were processed as described above to select the point mutant WT-mT/A. The mutant strain 3TER-mT/A was constructed in a similar way using the plasmid pBRL1162 as a template in the first PCR and WT-mT/A strain as a recipient for transformation. To construct the IPTG-controlled system for *rho* expression, the DNA fragment amplified from BSB1 chromosome by PCR with the oligonucleotides veb806 and veb880 was digested by EcoRI and BamHI endonucleases and clones at pMUTIN4 vector (Vagner et al., 1998). The resulting plasmid was transformed in BSB1 cells with selection to erythromycin resistance.

For the purification of the *B. subtilis* Rho protein, *rho* gene was amplified by PCR with oligonucleotides veb596 and veb599, treated by NdeI and SalI endonucleases and cloned at the expression vector pET28a (Novagen) allowing expression of Rho with N-terminal hexa-histidine tag (Ingham *et al*., 1999). The resulting plasmid pETRho was transferred to *E. coli* strain BL21-CodonPlus (DE3)-RIL (Stratagene).

### Luciferase assay

Analysis of promoters’ activity using luciferase fusions was performed as described previously (Mirouze *et al*., 2011) with minor modifications detailed in (Bidnenko et al., 2023). Cells were grown in LB medium to mid-exponential phase (OD_600_ 0,4-0,5), cultures were centrifuged and resuspended to OD 1.0 in fresh LB or DSM, to follow expression of the fusions during growth or sporulation, respectively. The pre-cultures were next diluted in respective media to OD_600_ 0.025. The starter cultures were distributed by 200µl in a 96-well black plate (Corning, USA) and Xenolight D-luciferin K-salt (Perkin, USA) was added to each well to a final concentration of 1.5 mg/ml. The cultures were grown under strong agitation at 37°C and analyzed in Synergy 2 Multi-mode microplate reader (BioTek Instruments). Relative Luminescence Units (RLU) and OD_600_ were measured at 5 min intervals. Each fusion-containing strain was analyzed at least three times. Each experiment included four independent cultures of each strain.

### Epifluorescence microscopy and image processing

For all microscopic observations, cells were mounted on a 2% agarose pad and topped with a coverslip. Bacteria were imaged with an inverted microscope (Nikon Ti-E) equipped with an iLas2 laser coupling system from Roper Scientific (150 mW, 488nm and 50 mW, 561 nm), a 100× oil immersion phase objective, an ORCA-R2 camera (Hamamatsu) and controlled by the MetaMorph software package (v 7.8; Molecular Devices, LLC). The post-acquisition treatment of the images was done with the Fiji software (van Ooij et al., 2004; Schindelin et al., 2012). To determine the frequency of cells entering into sporulation, cultures were sampled three hours after the induction of sporulation by the resuspension method (Harwood and Cutting,1990). Sampled cells were mixed with the lipophilic fluorescent dye Mitotracker Red (10 µg/ml final concentration) prior to microscopic observation. Asymmetric septa were manually counted in two independent replicas (N > 450 per strain and per replica).

To assess *rho* expression during sporulation using GFP as a reporter, cells bearing the non-modified or the point-mutated P*_rho_*-*gfp* transcriptional fusions were induced for sporulation by the resuspension method and sampled as above. Sampled cells were mixed with the lipophilic fluorescent dyes Nile Red (10 µg/ml final concentration) prior to microscopic observation. To measure the fluorescence intensity in the different compartments, circular areas of a constant 0.45 µm diameter were drawn in the center of individual compartments (forespores, mother cells, or predivisional cells) using images showing membrane-labelled cells, and recorded as a list of ROI. ROIs were subsequently applied over the corresponding image displaying the GFP signal, and the average fluorescence over each ROI recorded. Background fluorescence, determined as the average fluorescence from ROIs of identical areas spread over the field (outside cells), was subtracted to give the final fluorescence intensity of individual compartments. Counting was performed in at least two fields of view and for a minimum of 800 cell compartments per strain, in two independent experiments. Plotted values are average fluorescence intensities from pooled replicas, thus standard deviations account for cell to cell variability across all fields and replicas.

### RNA preparation and Northern blotting

Total RNA was extracted from *B. subtilis* cultures at OD_600_ indicated in the text by the glass beads method (Bechhofer et al., 2008). For Northern blotting, 5 µg of RNA were separated on 1% agarose-formaldehyde (2.2M) gel in MOPS (50mM)/EDTA (1mM) buffer (pH 7.0) and transferred to Hybond N+ membrane (GE-Healthcare) as described previously (Redko et al., 2013). Membrane pre-hybridization and the *rho* RNA detection by hybridization with the *rho*-specific α^32^P -labelled riboprobe was performed as detailed in (Gilet et al., 2020). The riboprobe was synthesized by T7 RNA polymerase in the presence of [α-^32^P]UTP at the template of the purified PCR fragment obtained with oligonucleotides YRH1 and YRH2 (S3 Table). The reverse PCR primer YRH1 contained at the 5’-end the T7 RNA polymerase promoter sequence (TAATACGACTCACTATA).

### Sporulation assay

Cells were induced for sporulation by the resuspension method, and the cultures were analyzed for the presence of spores starting from six hours after resuspension. The percentage of spores in a sample was calculated as proportion of viable cells after heating at 70°C for 15 min to the total number of cells as described in (Bidnenko et al., 2023).

### Spore purification and treatments

*B. subtilis* cells growing exponentially in LB were suspended in DSM at OD_600_ 0.05 and cultured at 37°C with aeration for 24 hours. Spores were purified by sequential rounds of intensive washing in ice-cold water and centrifugation over 3 days as described by Nicholson and Setlow in (Harwood and Cutting, 1990). The samples of the purified spores were controlled for the absence of cells by the heat-resistance test as described above and stored at 4°C in water at OD_600_ > 10. The purity of standard spore samples was above 95 percent.

For electron microscopy analysis, spores were additionally purified on Nicodenz (Axis-Shield, United Kingdom) density gradients. Spores were suspended in 1 ml of 20 % Nicodenz solution, layered on 50 % Nicodenz (15 ml) in the centrifuge tubes and centrifuged at 14,000 x g for 30 min at 10°C. The pelleted pure spores were washed 5 times in ice-cold water to remove traces of Nicodenz and kept at 4°C before analysis.

Prior the assays of spore resistance phenotypes and germination, the purified spores were suspended in 10 mM Tris HCl (pH 8.0) at OD_600_ 1.0 and activated by heating at 70°C for 30 min and subsequent cooling in ice for 20 min. The activated spores were used in the assays within 1 hour.

Chemical removal of spore coats was performed according to (Riesenman and Nicholson, 2000). Spores were suspended in decoating solution (50 mM Tris base, 8 M urea, 50 mM dithiothreitol, 1% sodium dodecyl sulfate; pH 10.0) at OD_600_ 5.0 and incubated at 60°C for 90 min with vortexing. After the treatment, spores were intensively washed in STE buffer (150 mM NaCl, 10 mM Tris-HCl, 1 mM EDTA; pH 8.0) as described.

### Transmission electron microscopy

Purified spores were fixed with 2% glutaraldehyde in 0.1 M sodium cacodylate buffer (pH 7.2) for 1 h at room temperature. Samples were contrasted with 0.5% Oolong tea extract in cacodylate buffer and postfixed with 1% osmium tetroxide containing 1.5% potassium cyanoferrate. The samples were dehydrated and embedded in Epon (Delta Microscopies, France), as described (Theodorou et al., 2019). Thin sections (70 nm) were collected onto 200-mesh copper grids and counterstained with lead citrate. Grids were examined with a Hitachi HT7700 electron microscope operated at 80 kV, and images were acquired with a charge-coupled device camera (Advanced Microcopy Techniques; facilities were located on the MIMA2 platform [INRAE, Jouy-en-Josas, France; https://doi.org/10.15454/1.5572348210007727E12]). The post-acquisition treatment of the images and measurement of the thickness of the spore coat were performed using the Fiji software (Schindelin et al., 2012).

### Spore UV resistance assay

The activated spores were suspended in Tris-HCl (pH 8.0) at OD_600_ 0.5, pipetted in triplicates (50µl) in the wells of 48-well sterile microtiter plates (Evergreen Scientific) and irradiated by ultraviolet (254 nm) light using Stratalinker 2400 UV Crosslinker (Stratagene) at the J/m^2^ doses indicated in the text (one plate per UV dose). The irradiated spore suspensions were plated in sequential dilutions at LB plates and the colonies were counted after 24 hours of incubation at 37 °C. The UV-resistance was determined as a ratio of the colony-forming units in the irradiated and non-irradiated samples of spores. Five experiments were performed with two independent sets of spores differentially expressing Rho.

To determine the effect of *rho* inactivation on the UV resistance of vegetative cells, WT and RM cultures in the late exponential phase (∼10^8^ cells/ml) were plated on LB plates and irradiated with the increasing doses of UV (254 nm) light using a Stratalinker 2400 UV Crosslinker (Stratagene). The UV-resistance of cells was estimated as above. Three independent experiments were performed.

### Dipicolinic acid (DPA) assay

The DPA content of spores was analyzed according to the protocol of Nicholson and Setlow (in Harwood and Cutting, 1990).

### Spore germination and outgrowth assay

Spore germination and outgrowth assays were performed in MS medium (10.8 g l^−1^ of K_2_HPO_4_, 6 g l^−1^ of KH_2_PO_4_, 1 g l^−1^ of C_6_H_5_Na_3_O_7_.2H_2_O, 0.2 g l^−1^ of MgSO_4_.7H2O, and 2 g l^−1^ of K_2_SO_4_) supplemented with 0.5% glucose, 0.01% L-tryptophan, 0.1% glutamate, 0.1 mM of FeCl_3_ citrate, 0.1 mM of CaCl_2_, 1 mM of MgSO_4_ and trace elements (0.001 g l^−1^ of MnCl_2_ 4H_2_O, 0.0017 g l^−1^ of ZnCl_2_, 0.00043 g l^−1^ of CuCl_2_· 2H_2_O, 0.0006 g l^−1^ of CoCl_2_ 6H_2_O and 0.0006 g l^−1^ of Na_2_MoO_4_· 2H_2_O). The stock solutions (100mM) of L-alanine and L-valine germinants were prepared in 10 mM Tris-HCl buffer (pH 8.0) containing 10mM D-glucose and 100mM KCl. The stock solution of AGFK germinant mixture contained 100 mM L-asparagine, 10mM D-glucose, 10 mM D-fructose and 100mM KCl in 10 mM Tris-HCl buffer (pH 8.0). The activated spores were diluted to OD_600_ 0.2 in the cold MS medium and distributed by 135 µl in the 96-well plate. Spore suspensions were simultaneously induced for germination by adding 15µl of a stock germinant solution, incubated under strong agitation at 37°C and monitored for OD_600_ at 2 min intervals in Synergy 2 Multi-mode microplate reader (BioTek Instruments). It usually took ∼1 min between germinant addition and the first OD_600_ reading. When needed, the stock solution of the germinant L-alanine was supplemented with 1mM IPTG or 5% casamino acids to get their final concentrations in spore suspensions 100µM and 0.5%, respectively. Three independently prepared sets of spores were used in the analysis. The assays were performed with each set of spores at least twice and included up to six replicas of each spore suspension in the same plate.

### Purification of the *B. subtilis* Rho protein for antibody preparation

*E. coli* BL21-CodonPlus (DE3)-RIL cells containing pETRho were grown to an OD_600_ of 0.2 in a 400 ml culture (2YT medium) at 16°C, and Rho expression was induced by the addition of 0.5 mM IPTG with continued growth overnight. The culture was harvested, pelleted and frozen at –80°C until further use. The frozen cells were resuspended in 10 ml of lysis buffer at 4°C containing 20 mM Tris–HCl pH 9.0, 100 mM Na_2_HPO_4_, 0.3 M NaCl, 10% glycerol and 0.1% Triton X-100, to which were added 10 mg/ml DNase I and an EDTA-free anti-protease tablet (Roche). The suspension was passed twice through a French press (20 000 p.s.i.) and the lysate centrifuged for 30 min at 15 000 *g*. Imidazole-HCl (pH 8.0) was added to the supernatant to give a final concentration of 1 mM and the resultant suspension was applied to a 1 ml Ni^2+^-NTA column (Qiagen). The resin was then washed sequentially with 10 ml of lysis buffer (without Triton X-100), 10 ml of buffer containing 20 mM Tris–HCl pH 9.0, 0.3 M NaCl, 20 mM Imidazole and, finally, 10 ml 20 mM Tris–HCl pH 9.0, 0.3 M NaCl and 250 mM Imidazole. *B. subtilis* Rho was eluted with 20 mM Tris–HCl pH 9.0, 100 mM Na_2_HPO_4,_ 0.3 M NaCl and 500 mM Imidazole as 1ml fractions into collection tubes pre-filled with 3 ml elution buffer lacking Imidazole, to immediately dilute the sample 4-fold and avoid precipitation. The protein peak was determined initially by measuring the protein concentration (Bio-Rad) in the different fractions. Peak fractions were pooled and dialyzed over-night in buffer containing 20 mM Tris–HCl pH 9.0, 100 mM Na_2_HPO_4_, 0.3 M NaCl, 10% glycerol. Protein purity was verified by SDS–PAGE analysis and estimated at >95%. A sample of purified *B. subtilis* Rho protein at 1.1 mg/ml in storage buffer was filtered through 5µm, 0.5µm and then 0.2µm Acrodisk filters before rabbit immunization to generate custom anti-Rho polyclonal antibodies through the commercial ‘87-day anti-antigen classical’ program of Eurogentec (Belgium). The low solubility of *B. subtilis* Rho in buffers suitable for immobilization on affinity chromatography columns prevented further purification of the anti-Rho antibodies from the crude sera. The best anti-Rho serum aliquot was selected from the immunization program aliquots by Western blotting with purified *B. subtilis* Rho and *B. subtilis* cell extract samples.

### Western blotting

The crude cell extracts were prepared using Vibracell 72408 sonicator (Bioblock scientific). Bradford assay was used to determine total protein concentration in each extract. Equal amounts of total proteins were separated by SDS-PAGE (10% polyacrylamide). After the run, proteins were transferred to Hybond PVDF membrane (GE Healthcare Amersham, Germany), and the transfer quality was evaluated by staining the membrane with Ponceau S (Sigma-Aldrich). The Rho protein was visualized by hybridization with antiserum against *B. subtilis* Rho (Eurogentec, Belgium; dilution 1:5,000) and the secondary peroxidase-coupled anti-rabbit IgG antibodies A0545 (Sigma-Aldrich; dilution 1:20,000).

## Supporting information

**S1 Fig.**
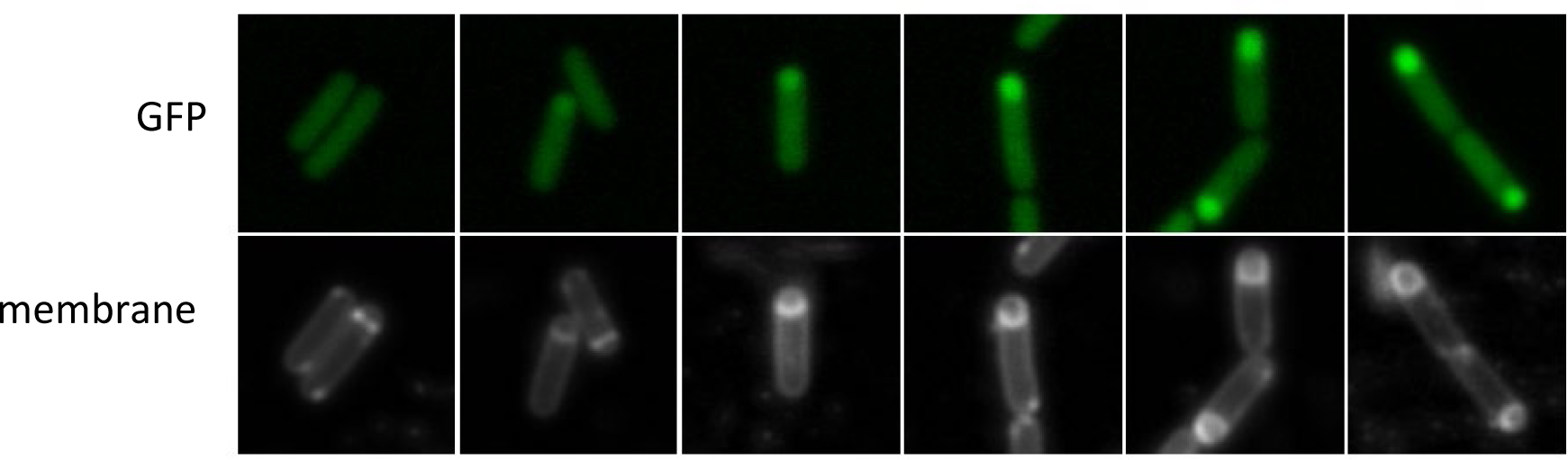
Induction of P*rho*-*gfp* expression in the forespore follows the progression of the engulfment process. Montage of fluorescent microscopy images (upper row: green channel; lower row: red channel) of individual wild type cells bearing P*_rho_*-*gfp* transcriptional fusion during the growth in DSM. Cells were labeled with the red fluorescent Nile-Red dye and observed 2 hours after entering stationary phase (T2). Images are displayed from left to right following the progression of the sporulation process, from predivisional to fully engulfed prespore state (red channel), to show an increase of the fluorecence levels (green channel) in the prespore.

**S2 Fig.**
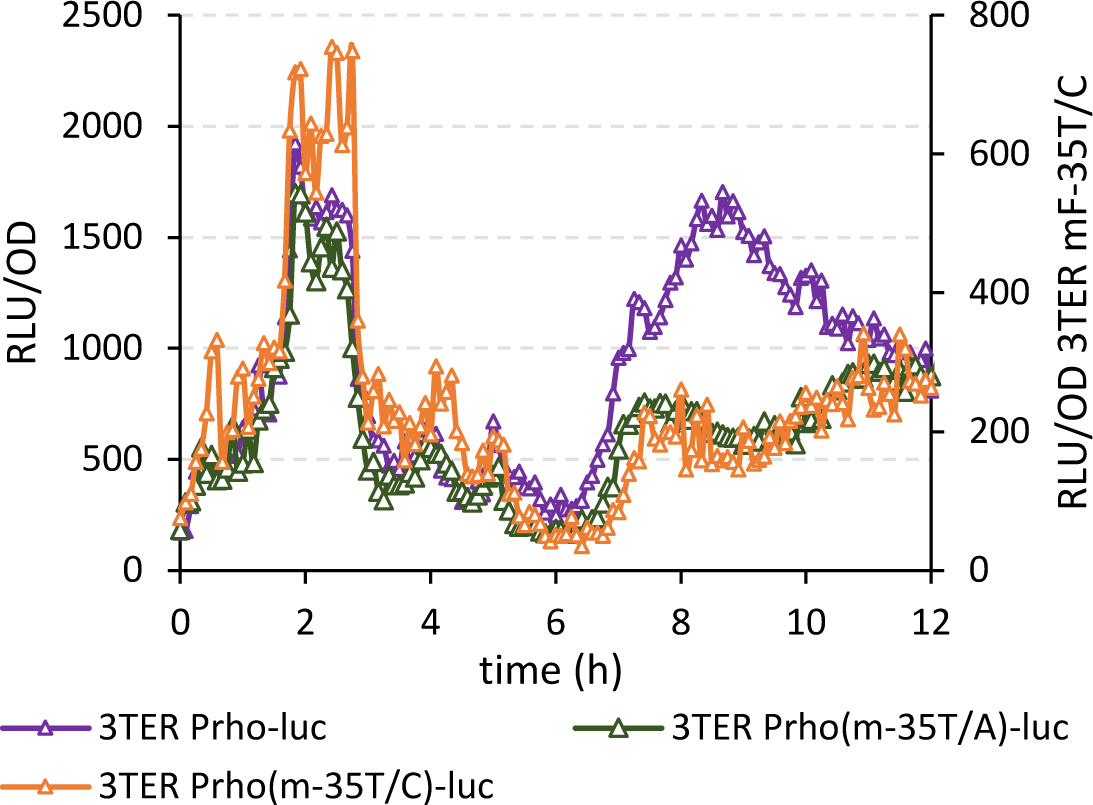
Inactivation of the SigF-dependent *rho* promoter inhibits sporulation-specific expression of luciferase in 3TER P*_rho_-luc* cells. The mutational analysis of the SigF-dependent promoter of *rho* using luciferase was performed in the 3TER genetic background to limit the interference of the read-through transcription. 3TER cells bearing a non-modified (violet triangles) or the point-mutated *mF-35T/A* (green triangles) and *mF-35T/C* (orange triangles) P*_rho_-luc* fusions were set to sporulate in DSM and analyzed for luciferase activity as described in Fig 2. Of note, the mutation *mF-35T/A* specifically decreases luciferase expression during sporulation, while the mutation *mF-35T/C* has a global inhibitory effect (scaled differently in the graph). Nevertheless, the sporulation-specific luminescence in 3TER *mF-35T/C* P*_rho_-luc* is affected to a similar extent as in 3TER *mF-35T/A* P*_rho_-luc*.

**S3 Fig.**
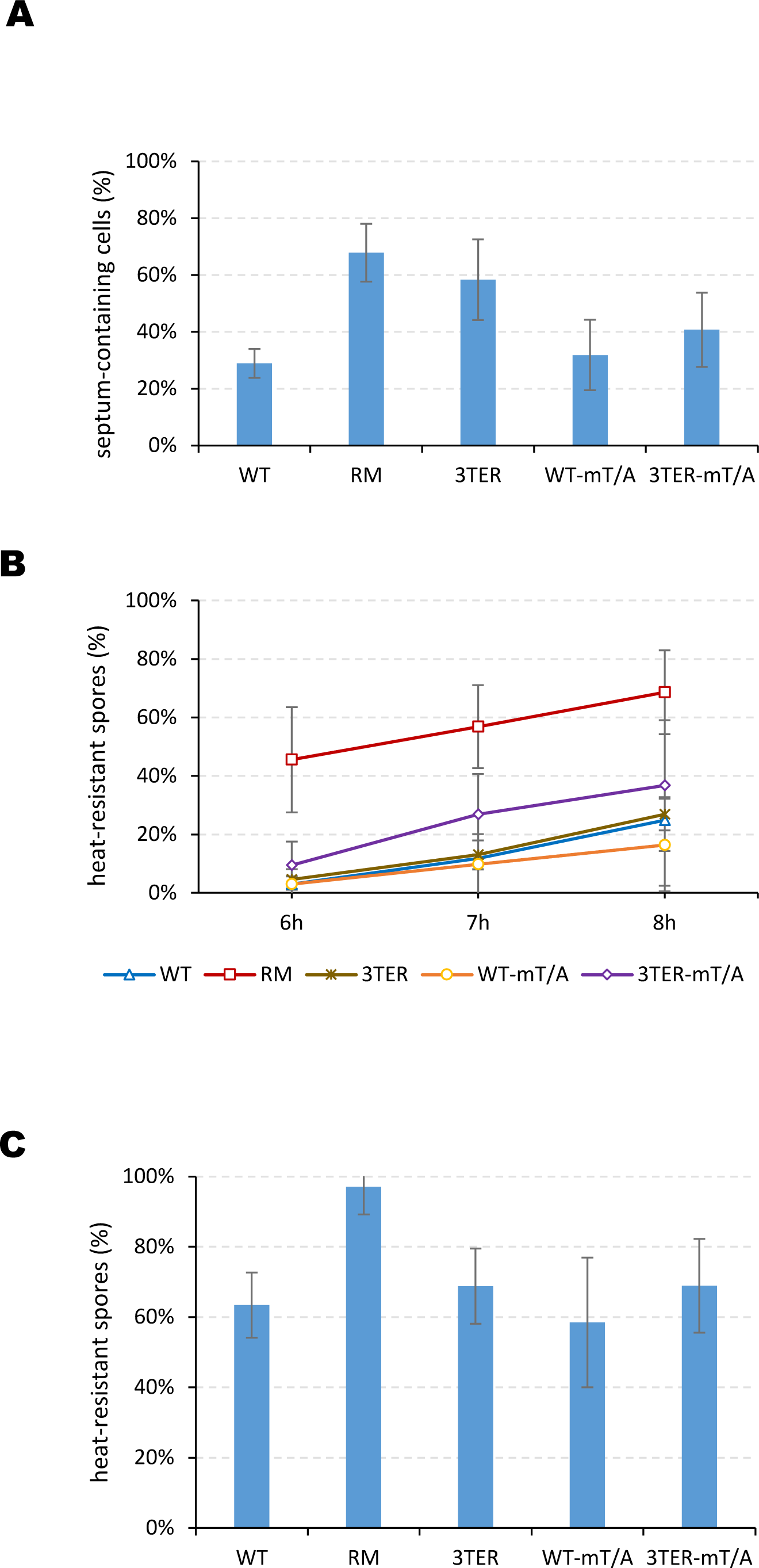
Complete inactivation of *rho* has a maximal effect on sporulation efficiency. (A) Cells were induced for sporulation by the resuspension method, sampled three hours after resuspension, colored with lipid affine Mito-tracker red and analyzed by fluorescence microscopy for the presence of asymmetric septum. The proportion of septum-containing cells was determined by manual counting, in at least three fields of view and for a minimum of 600 cells per strain and per replica. Plotted are average proportions of septum-containing cells from three experimental replica. The bars represent standard deviation from the mean values. (B and C) After resuspension, cells were continuously propagated in a poor SM medium and analyzed for the formation of heat-resistant spores at the indicated time (B) and overnight (20 hours) (C), as described in Materials and Methods. Plotted are average proportions of the heat-resistant spores from three independent experiments, two of which were performed in a prolongation of microscopy analyses in (A). The bars represent standard deviation from the mean values.

**S4 Fig.**
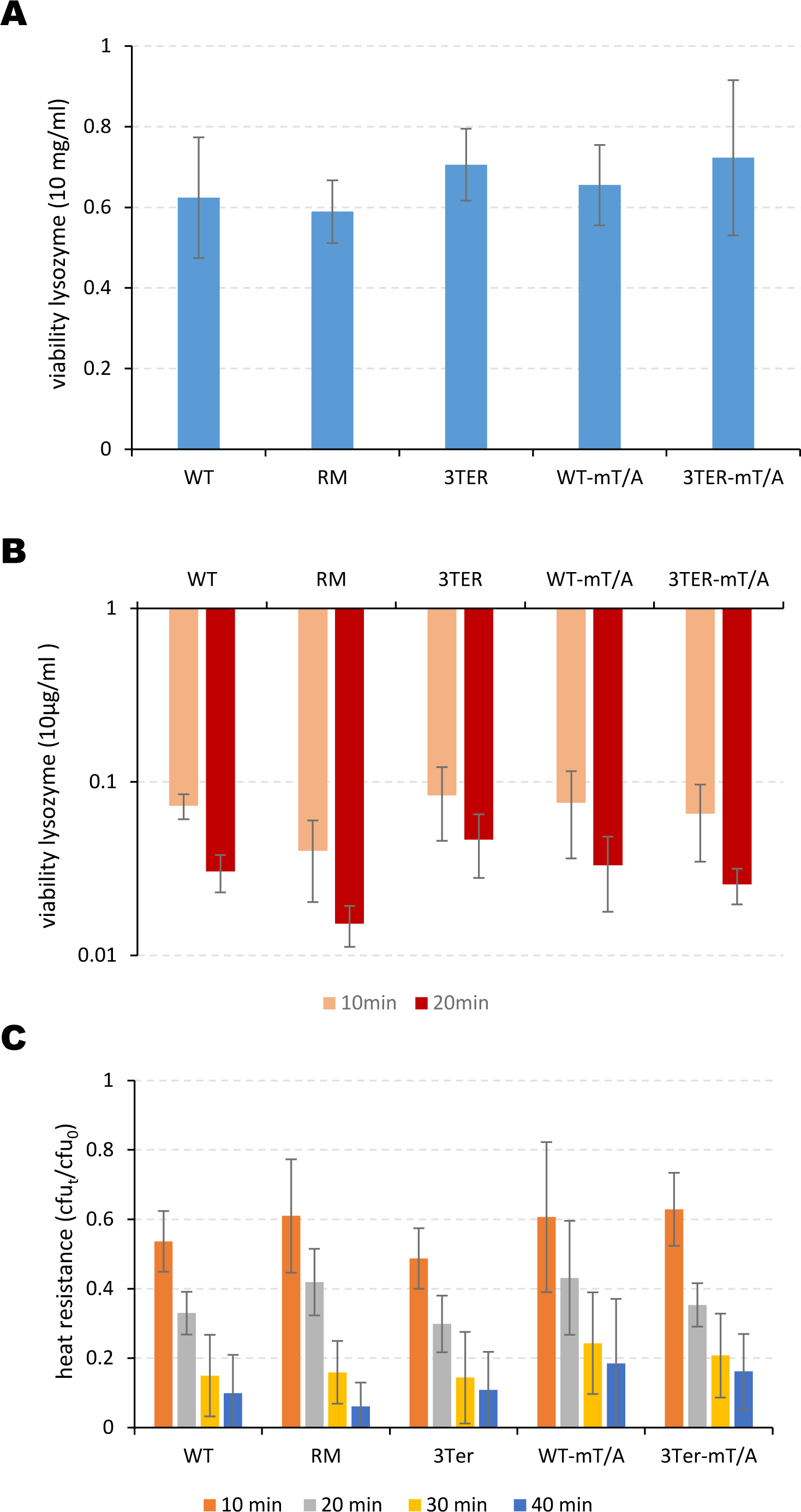
Alteration of *rho* expression does not significantly affect the resistance of spores to heat and lysozyme. The purified spores of the wild type strain (WT), *rho*-deletion mutant (RM) and strains altered for *rho* expression in the mother cell (3TER) or in the forespore (WT-mT/A) or in the two compartments of sporangia (3TER-mT/A) were analyzed for the resistance to lysozyme (A and B) and heat (C). (A) Spores were activated as described in Materials and Methods, suspended in TE buffer (10mM Tris-HCl, 1mM EDTA, pH 8.0) containing 10 mg/ml lysozyme (Fluka) at OD_600_ 0.2 and incubated at 37°C for 1 hour. (B) Spores were chemically treated to remove spore coats as described in Materials and Methods. Decoated spores were suspended in 10mM Tris-HCl (pH 8.0) at OD_600_ 0.2 and treated with lysozyme (10 µg/ml, final concentration) for 10 and 20 minutes at 37°C. In (A) and (B), spore viability after lysozyme treatment was established by plating spores from the treated and untreated samples at LB agar plates. (C) The activated spores were heated at 90°C for the indicated time, platted at LB agar plates and incubated at 37°C for 24 hours. The heat resistance of spores was calculated as a proportion of viable colony-forming spores in heated and unheated samples. The same set of spores was used in all experiments; spore samples in (B) originate from two independent decoating procedures. The experiments were reproduced four times, and the established average values are presented with the standard deviations of the mean (bars).

**S5 Fig.**
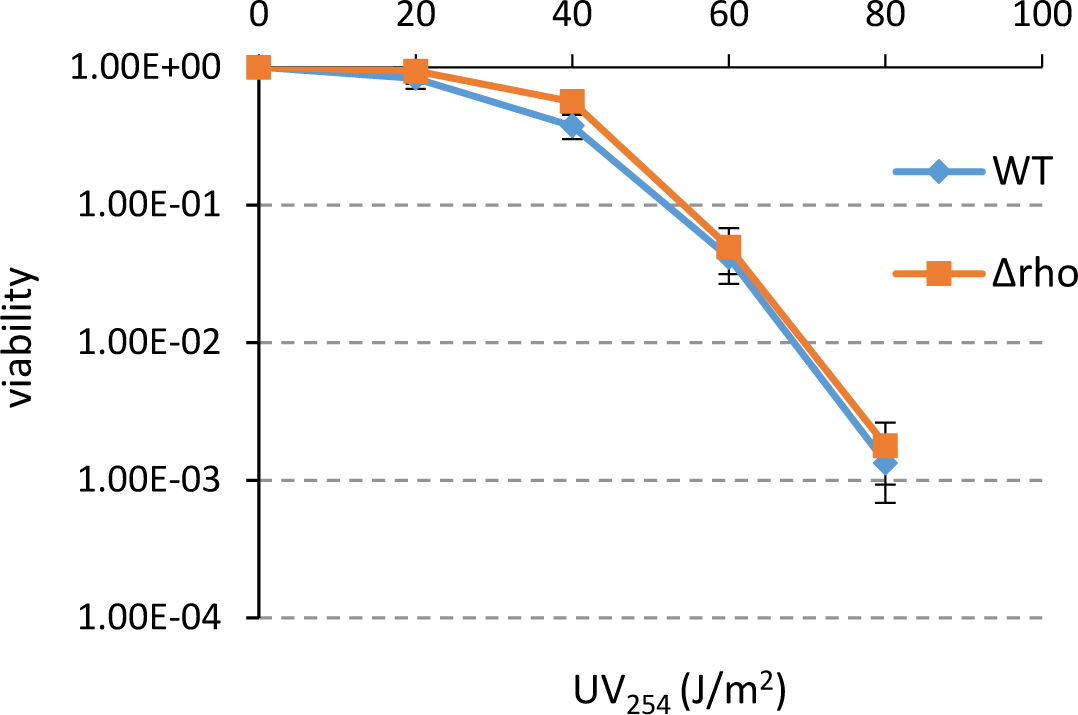
Rho inactivation does not affect UV-resistance of vegetative cells. The wild type (WT, blue line) and Δ*rho* (RM, red line) cells were grown in LB to OD_600_ 0.5, platted at LB-agar plates in serial dilutions and irradiated by ultraviolet light at the indicated doses using the UV crosslinker as described in Materials and Methods. Cells were scored after 24h of incubation at 37°C. The experiment was reproduced twice with three culture samples of each strain. Plotted are the mean values from two experiments.

**S6 Fig.**
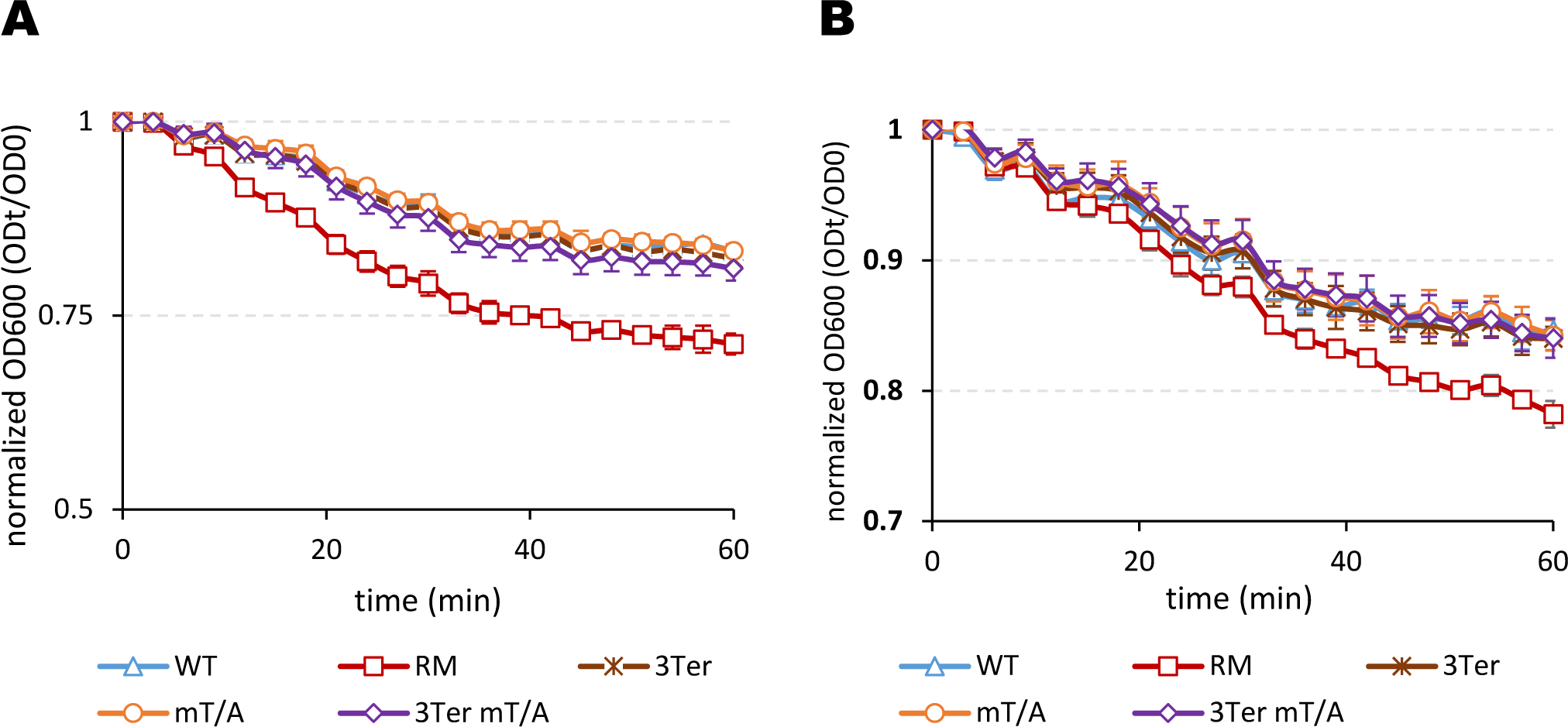
Absence of Rho accelerates spore germination with different nutrient germinators. Spores produced by WT (blue triangles), RM (red squares), 3TER (brown crosses), WT-mT/A (orange circles) and 3TER-mT/A (violet diamonds) cells were induced for germination by (A) the germinant AGFK (10 mM L-asparagine, 1mM D-glucose, 10mM KCl and 10 mM D-fructose, final concentrations) or (B) L-valine (10mM). The experiments were performed twice with two independent sets of spores. Each experiment included up to six replicas of individual suspensions of spores. The results of the representative experiment are plotted. The bars represent standard deviation of the mean values.

**S7 Fig.**
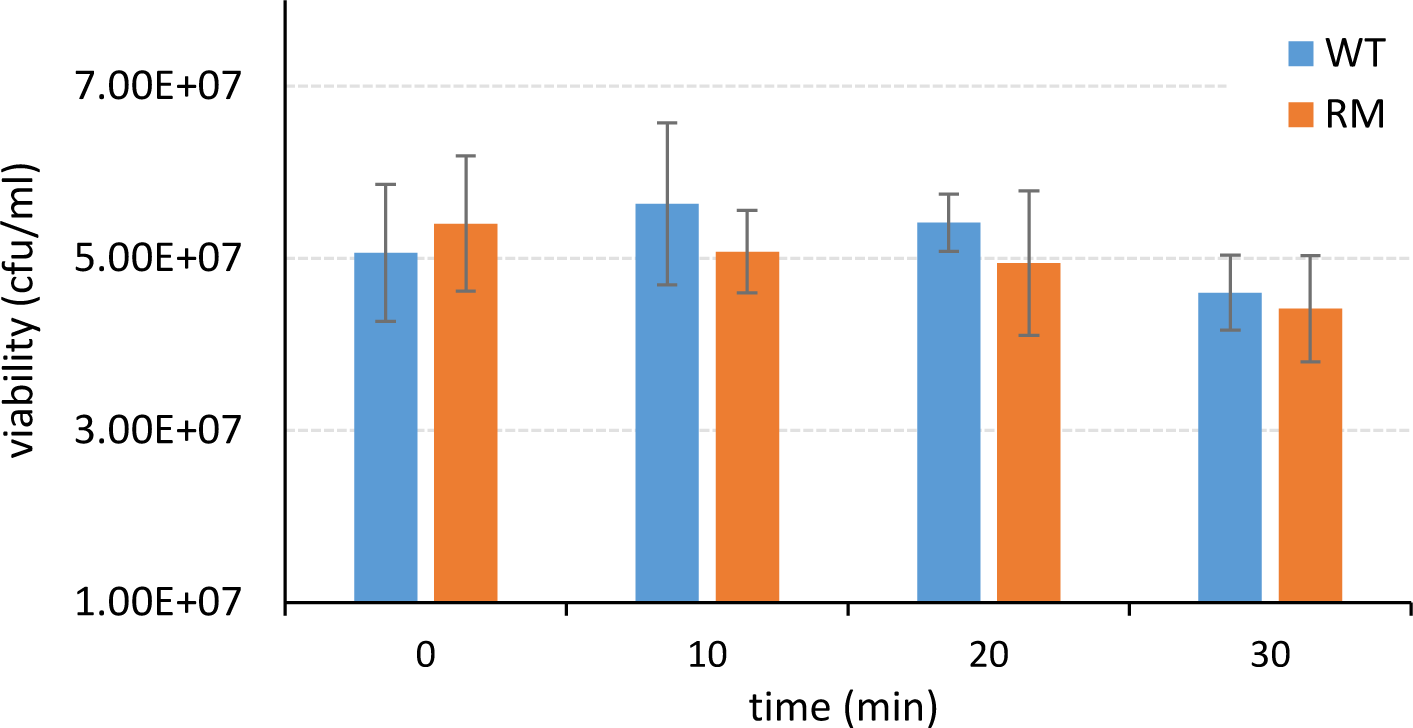
Absence of Rho does not affect viability of germinating spores. The wild type WT (blue) and *rho*-minus RM (red) spore suspensions at OD_600_ 0.1 were activated and induced for germination by 10mM L-alanine as described in Materials and Methods. Spore viability during germination was established by platting of spores in sequential dilutions at LB-agar plates at the indicated time after germinant addition. Plates were incubated at 37°C for 24h before the colonies counting. The experiment was reproduced twice with two independently prepared spore samples and included four replicas of each spore suspension. The results of the representative experiment are plotted. The bars represent standard deviation from the mean values.

**S8 Fig.**
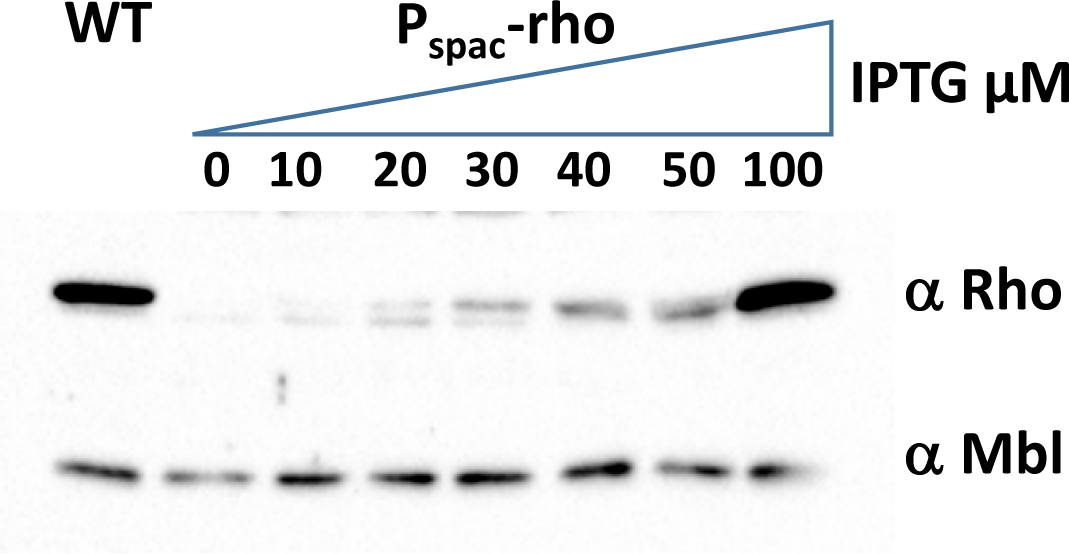
Functional analysis of P*_spac_*-*rho* transcriptional fusion. *B. subtilis* cells expressing *rho* from the IPTG-inducible promoter (P_spac_-*rho*) were incubated in LB medium in the presence of the indicated concentrations of IPTG to OD_600_ 0.6 and analyzed for the levels of Rho protein in comparison with wild-type cells (WT) by Western blot with the specific anti-Rho^Bs^ anti-serum as described in Materials and Methods. Protein equilibrium between the loaded samples was controlled by Bradford assay and visualization of Mbl protein by anti-Mbl antibodies as described in (Bidnenko et al., 2023).

**S1 Table. Differential expression analysis of genes from the “Sporulation” lifestyle category in stationary *Δrho* vs. stationary WT *B. subtilis* cells (compiled from S3 Table, Bidnenko et al., 2023).**

**S2 Table.**
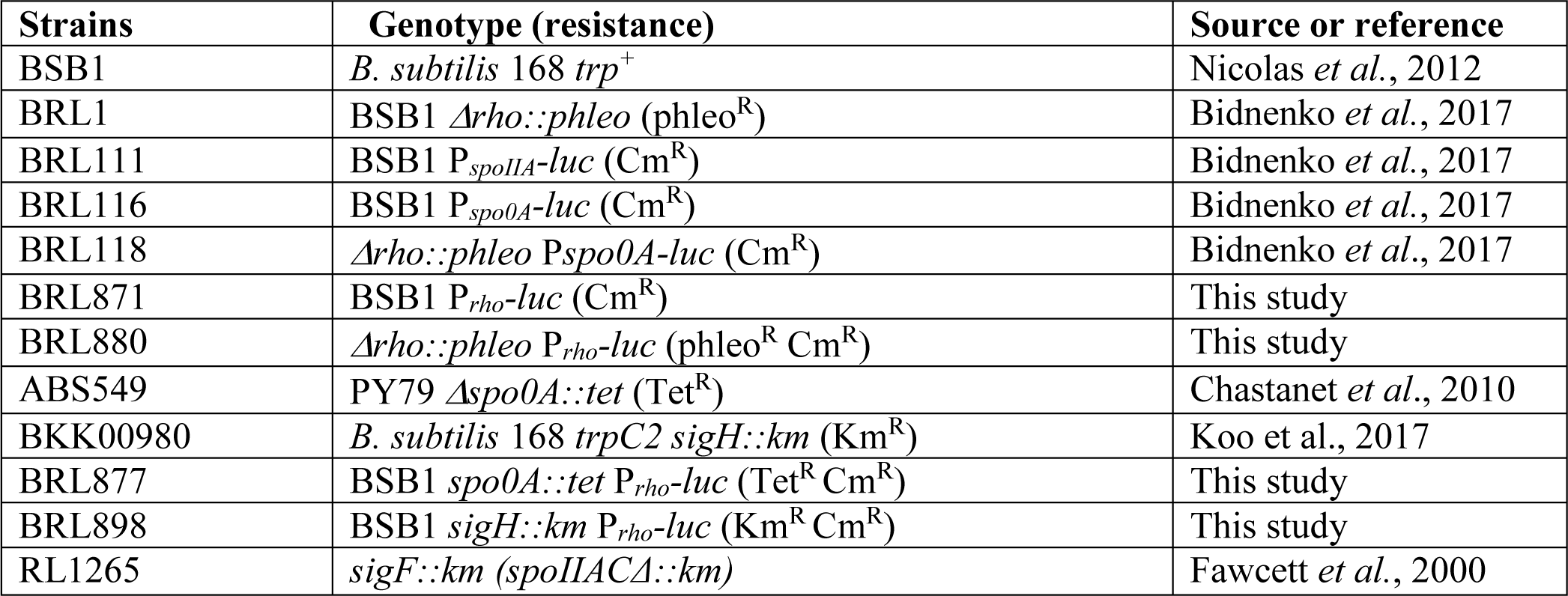
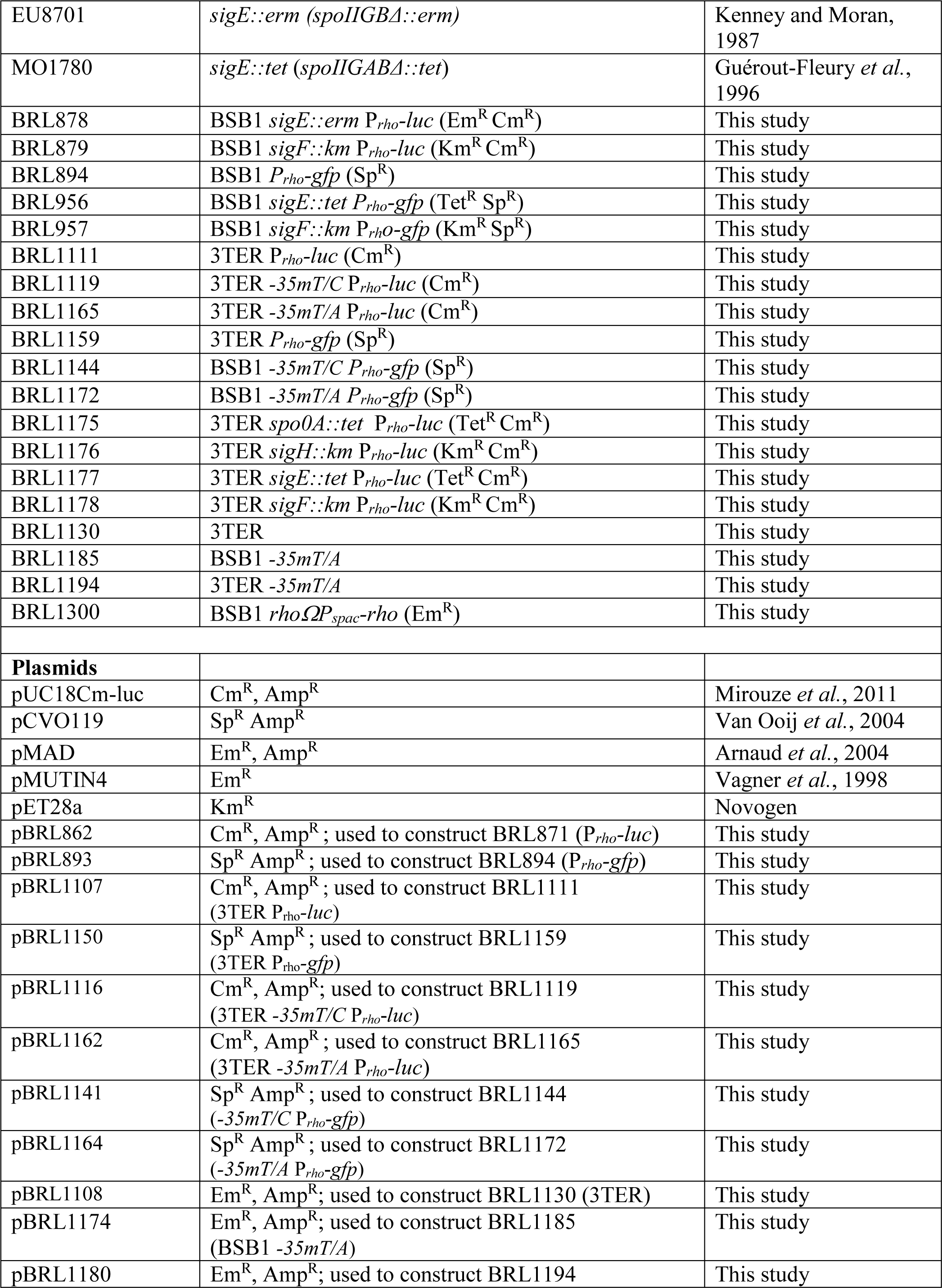
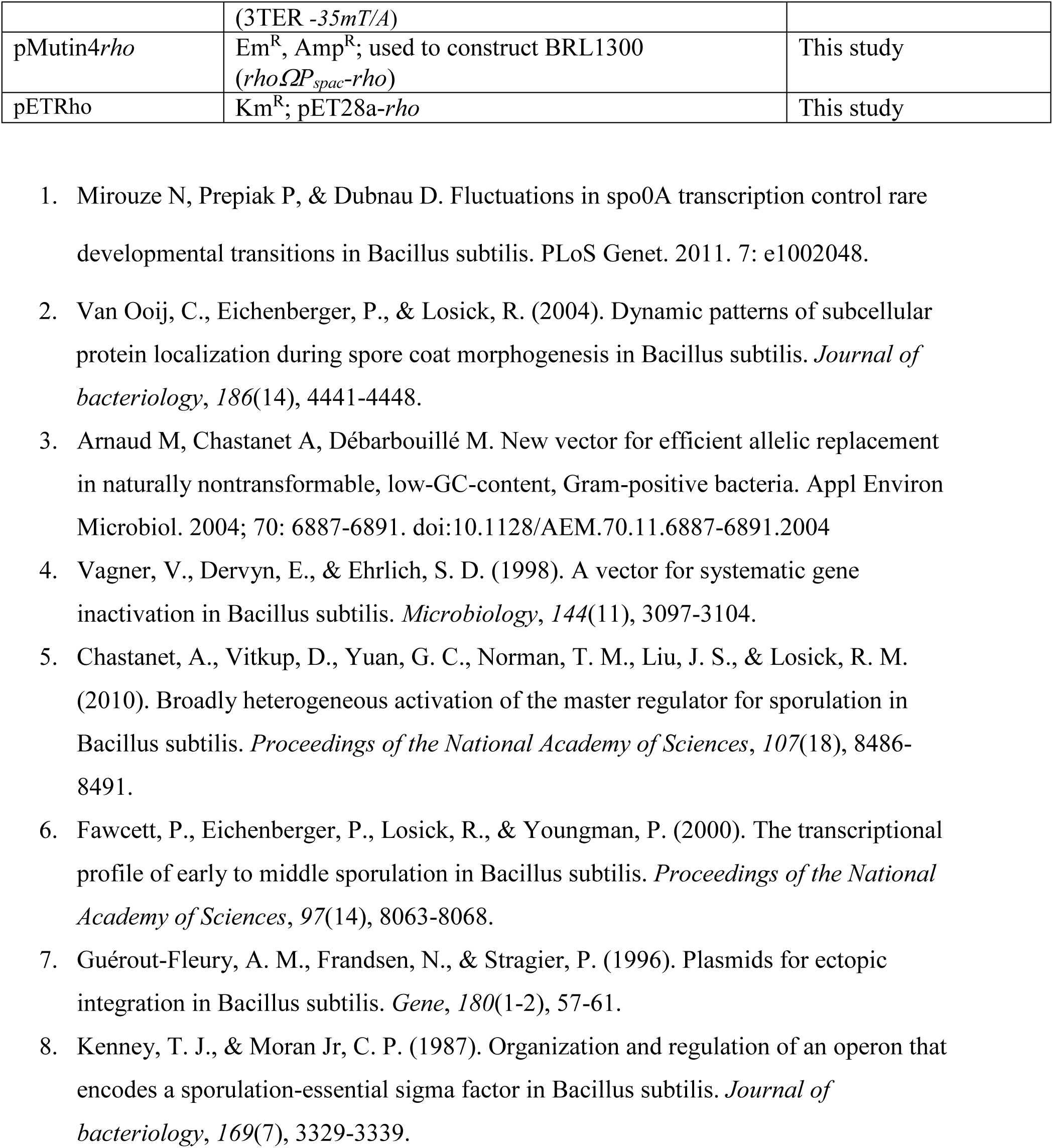
Strains and plasmids used in the analysis.

**S3 Table.**
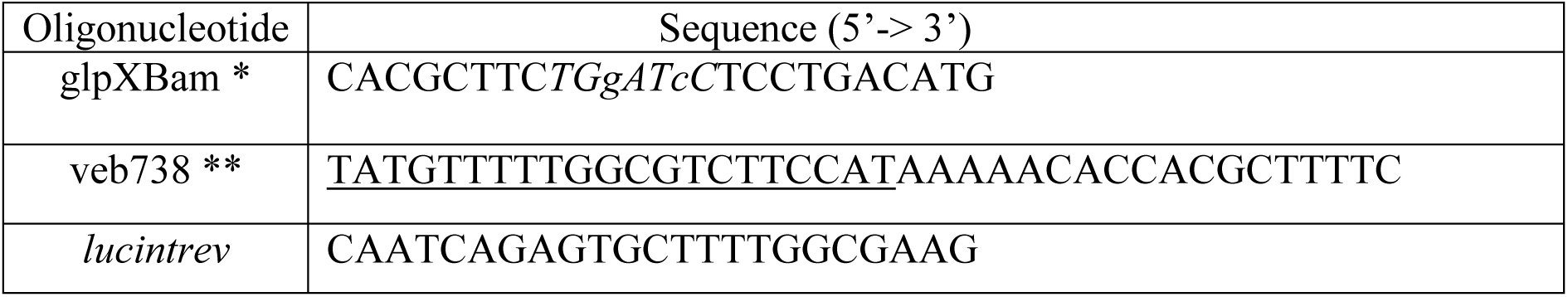
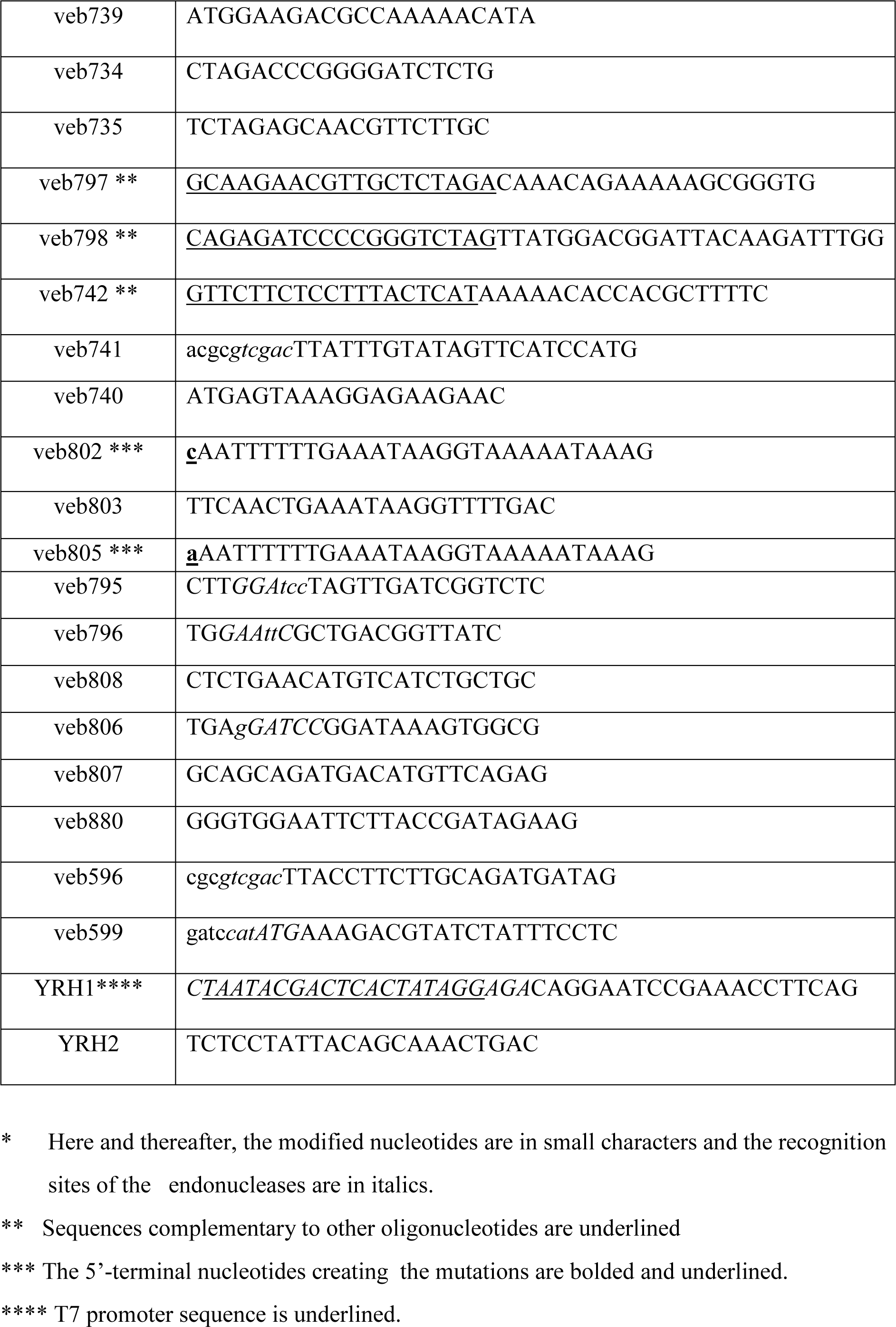
Oligonucleotides used for strains construction.

